# Gene synteny and translational coupling of *sctS* and *sctT* facilitate assembly of the unique helical T3SS export apparatus in *Salmonella* Typhimurium

**DOI:** 10.64898/2026.02.19.706803

**Authors:** Eunjin Kim, Mirjam Forberger, Felix Weichel, Claudia Paroll, Jialin Zhou, Iwan Grin, Samuel Wagner

**Affiliations:** University of Tübingen, Interfaculty Institute of Microbiology and Infection Medicine (IMIT), Section of Cellular and Molecular Microbiology, Elfriede-Aulhorn-Str. 6, 72076 Tübingen, Germany; German Center for Infection Research (DZIF), partner-site Tübingen, Elfriede-Aulhorn-Str. 6, 72076 Tübingen, Germany; Excellence Cluster “Controlling Microbes to Fight Infections” (CMFI), Elfriede-Aulhorn-Str. 6, 72076 Tübingen, Germany

## Abstract

Virulence-associated type III secretion systems (T3SS) are employed by many gram-negative pathogens to translocate effector proteins into hosts. These nanomachines are composed of cytoplasmic components, a needle complex base, housing the export apparatus and anchoring the machine in the bacterial cell envelope, a needle filament, and a translocon that penetrates the host membrane. Assembly of the T3SS is a hierarchical process, which is initiated by coordinated association of the five subunits comprising the export apparatus, SctR, SctS, SctT, SctU, and SctV, in the bacterial inner membrane. In this study, we report that proper assembly of this uniquely helical export apparatus is fine-tuned by the genetic organization of the export apparatus genes in *Salmonella* Typhimurium. The *sctS* mRNA harbors a stem-loop structure which conceals the Shine-Dalgarno sequence of *sctT* from recognition by a ribosome. *sctT* translation is only possible upon melting of the stem-loop by a *sctS*-translating ribosome. This mechanism translationally couples *sctS* and *sctT.* We show that this strict regulation prevents uncontrolled overexpression of SctT and its assembly into futile multimers that disrupt export apparatus assembly and reduce pathogen fitness. Based on the strong synteny of T3SS export apparatus genes and their highly conserved unique structure, also among closely related bacterial flagella, similar mechanisms are likely needed to enable a tightly regulated and stoichiometric assembly process of these molecular machines in general.

## Introduction

Virulence-associated type III secretion systems (T3SS), whose core structure is also known as injectisome, are a highly conserved protein complex mediating translocation of a variety of effector proteins into host cells. T3SSs are large machines with a size of ∼7 MDa, consisting of about 200 subunits and spanning the bacterial inner (IM) and outer (OM) membranes as well as a host membrane (Zilkenat et al., 2016). These nanomachines consist of several substructures, including the export apparatus, the membrane-anchored base, the cytoplasmic components, and the needle and tip.

The export apparatus is a core component of the T3SS, localized within the membrane-anchored base. The export apparatus, which is also conserved in the closely related bacterial flagella, is composed of five subunits, namely SctR, SctS, SctT, SctU, and SctV, with a stoichiometry of 5:4:1:1:9 (Kuhlen et al., 2018; Zilkenat et al., 2016). SctRSTU constitute the core export apparatus, which forms a pore that T3S substrates pass through before entering the secretion channel formed by needle adapter and filament proteins (Hu et al., 2019; Torres-Vargas et al., 2019). In non-secreting conditions, the export apparatus pore is gated by the SctT plug, but contact with substrates results in opening of the gate and substrate secretion (Hüsing et al., 2021; Kuhlen et al., 2018; Miletic et al., 2021). The export apparatus has been shown to contribute to hierarchical secretion of substrates through SctU (Björnfot et al., 2009; Edqvist et al., 2003; Inoue et al., 2019; Monjarás Feria et al., 2015) and SctV (Marcos-Vilchis et al., 2025; Portaliou et al., 2017; Yuan et al., 2021). SctV directly interacts with the substrates and their chaperones through its cytoplasmic domain (Gilzer et al., 2022; Portaliou et al., 2017; Xing et al., 2018) and also couples the proton motive force with substrate translocation (Erhardt et al., 2017; Minamino et al., 2011, 2016).

Recent structural studies of the export apparatus have shed light on its unique structural features. The core export apparatus (SctR_5_S_4_T_1_U_1_) is a helical pore, resembling an inverted cone, with the tip oriented toward the cytoplasmic side. Although the subunits are predicted to be transmembrane proteins, the core export apparatus is not fully integrated into the IM but instead protrudes significantly into the periplasmic side (Kuhlen et al., 2018). The core export apparatus displays pseudo-hexameric symmetry, which results from structural similarity between SctT and the combination of SctR and SctS (Kuhlen et al., 2018). In addition, the export apparatus exhibits a right-handed helical structure, characterized by the hydrophobic alpha helices being staggered by approximately 25 Å. The staggered conformation is stabilized by inter- and intramolecular salt bridges (Kuhlen et al., 2018; N. Singh et al., 2021). This helical configuration templates the helical needle adapter and filament in the downstream T3SS assembly (Goessweiner-Mohr et al., 2019; Torres-Vargas et al., 2019).

Based on biochemical and structural data, the helical assembly of the export apparatus is believed to occur in a sequential manner (Dietsche et al., 2016; Johnson et al., 2019; Kuhlen et al., 2018; N. Singh et al., 2021). Assembly begins with the formation of pentameric SctR, followed by the integration of one SctT molecule into the SctR pentamer to form a stable SctR_5_T_1_ core, which serves as a pseudo-hexameric building block. Four SctS peripherally assemble onto the SctR_5_T_1_ core in a sequential manner, forming SctR_5_S_4_T_1_ (Dietsche et al., 2016; Johnson et al., 2019; Kuhlen et al., 2018; N. Singh et al., 2021). The assembly of the core export apparatus is completed by recruitment of one SctU (Kuhlen et al., 2020; Wagner et al., 2010). Once the assembly of the nonameric SctV is completed, the core export apparatus is repositioned from the IM into the periplasmic space, a process that is further stabilized by interactions with the IM base substructures (Kuhlen et al., 2018). The export apparatus has been shown to be a critical starting point for the T3SS assembly process (Diepold et al., 2010; Wagner et al., 2010). It facilitates the assembly of the base as a scaffold, which in turn allows the assembly of the cytoplasmic components, enabling secretion and assembly of the needle components (Wagner et al., 2010).

As the nucleus and scaffold of T3SS assembly, the export apparatus must be efficiently and correctly assembled. While many protein complexes rely on diverse regulatory mechanisms to achieve stoichiometric and orderly assembly rather than relying on stochastic interactions, the regulation of export apparatus assembly remains poorly understood. Previous studies have shown that the core export apparatus genes exhibit high synteny (Abby & Rocha, 2012), which indicates physiological relevance, and often contain regulatory motifs. In the context of an operon, gene order dictates the assembly order of the protein complex, especially when the order is conserved (Wells et al., 2016). However, in the case of the export apparatus, gene order does not reflect assembly order. This observation led us to investigate whether the highly conserved gene order plays a role in regulating the assembly of this unique helical export apparatus.

We demonstrate that the conserved gene order of the export apparatus harbors a secondary RNA structure within the *sctS* mRNA, which tightly regulates the translation of *sctT* in *Salmonella* Typhimurium. This mechanism enables translational coupling between *sctS* and *sctT*, ensuring precise stoichiometric control. This regulation is particularly crucial for SctT, as SctT is overexpressed in the absence of the regulation and self-assembles into cytotoxic, unregulated multimers that may form leaky pores in the bacterial inner membrane. Thus, the translational coupling between *sctS* and *sctT* plays a vital role in maintaining robust and efficient assembly of the export apparatus and thus vitality of the bacteria harboring T3SS.

## Results

### Genes of the T3SS core export apparatus show a strict synteny

It was noted that the export apparatus genes are conserved in their genomic order. This drew our interest, especially given that the assembly order of the export apparatus differs from the order of the respective genes (Fig. 1A and 1B). In order to analyze the conservation of the gene order of the export apparatus components systematically, we performed a comprehensive *in silico* analysis of its gene organization in many bacterial species expressing T3SS.

**Figure 1.**
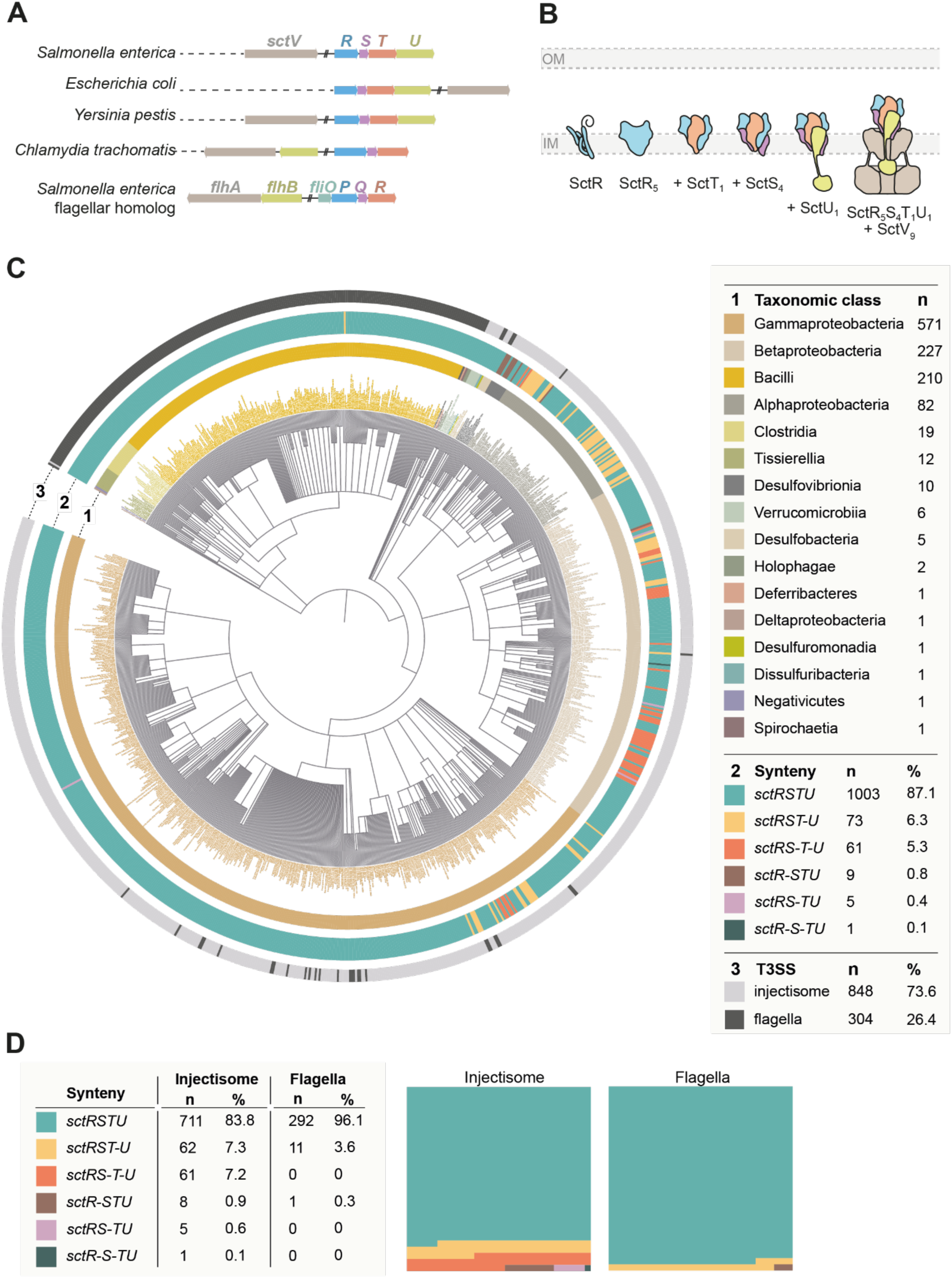
Gene order conservation of the T3SS export apparatus. **A.** Overview of the genetic organization of the genes of the T3SS export apparatus in selected organisms. Genes are color coded according to homology. **B.** Assembly of the T3SS export apparatus. The export apparatus assembles in the bacterial inner membrane (IM), starting with the SctR pentamer assembly. Subcomplexes with SctT, SctS, SctU, and SctV subunits are sequentially assembled in the indicated order. The export apparatus then serves as a nucleus for downstream T3SS assembly. IM, inner membrane. Adapted from (Pais et al., 2023). **C.** *In silico* analysis of the genetic organization of the export apparatus was performed using the Cblaster program (Gilchrist et al., 2021), with export apparatus genes from *S*. Typhimurium flagella (strain LT2) as query sequences. Among the Cblaster cluster hits, NCBI reference genomes containing all *sctRSTU* and the homologous *fliPQRS* genes were included for synteny analysis (n = 1152). Genes separated by more than 500 bp or located on different strands were considered disconnected (indicated by “–”). The taxonomic distribution of export apparatus gene clusters was mapped and visualized as a circular tree: ring 1 indicates the taxonomic class, ring 2 the synteny category, and ring 3 the T3SS type (injectisome vs. flagellum), with all categories color-coded as in the legend. **D.** Synteny distributions from panel C were quantified separately for injectisome and flagellar export apparatus gene clusters. The panels on the right show part-to-whole graphs for each system.

Using the Cblaster program (Gilchrist et al., 2021), we performed BLAST searches to identify homologs of the *sctRSTU* gene cluster. To minimize redundancy, particularly from overrepresented or extensively sequenced strains, only hits from bacterial reference genomes available in NCBI were included. Genomes containing all four *sctRSTU* genes were retained for further synteny analysis, which resulted in 1,152 bacterial genomes. These hits were categorized into different groups, based on each gene’s strand (+ or -) and locus information, and the intergenic distances between the genes in a cluster. Only when all genes in a *sctRSTU* cluster were localized to either the positive or negative strand and had intergenic distances of less than 500 bp, the hits were considered as having the synteny. Among the 1,152 export apparatus clusters, 848 were of T3SS (injectisome) and 304 were of flagella (Fig. 1C). In both injectisome and flagellar export apparatus genes, *sctRSTU* genetic organization was highly conserved, being present in 87.1 % (n = 1,003) in all analyzed genomes (83.8 % and 96.1 % in export apparatus gene clusters of injectisome and flagella, respectively) (Fig. 1C and Supplementary Table 2). In 3.6 % of flagella (n = 11), the *sctU* gene was separated from *sctRST* (*sctRST-U*), which is the case for *S*. Typhimurium strain LT2 genome. In injectisomes, this was7.3 % (n = 62). Other genetic organizations were additionally observed in injectisome export apparatus clusters, including *sctRS-T-U* (7.2 %), *sctR-STU* (0.9 %), *sctRS-TU* (0.6 %), and *sctR-S-TU* (0.1 %). The *sctRS-T-U* was predominantly observed in the *Burkholderiaceae* family, whereas the *sctRST-U* was mainly found in the *Alphaproteobacteria* class (Figure 1C and 1D, and Supplementary Table 2).

### Effective plasmid-based complementation of a chromosomal *sctT* deletion mutant requires expression of more than just *sctT*

We then sought to investigate the functional implications of the observed high level of gene order conservation using the export apparatus of the T3SS encoded by *Salmonella* pathogenicity island 1 (SPI-1) as a model. We reasoned that *in trans* complementation may be compromising if the synteny of export apparatus genes is important for assembly.

*S*. Typhimurium strains with individual deletions of export apparatus genes (Δ*sctR,* Δ*sctS,* Δ*sctT,* Δ*sctU*) were complemented *in trans* from rhamnose-inducible pT10 plasmids expressing a combination of export apparatus subunits. The secretion functionality of the assembled T3SS was assessed in each strain by luminescence measurements of secreted SipA-NanoLuc (NL) fusion protein (Fig. 2A) (Westerhausen et al., 2020). Δ*sctR,* Δ*sctS,* and Δ*sctU* strains exhibited comparable secretion levels of 50% of the wild-type when the missing export apparatus subunit was expressed alone or together with other subunits. In contrast, complementation of the Δ*sctT* mutant only reached secretion of less than 20% of the wild-type except when all four export apparatus genes were expressed together. These results indicate that the genetic context of *sctT* is critical for the assembly of the export apparatus.

**Figure 2.**
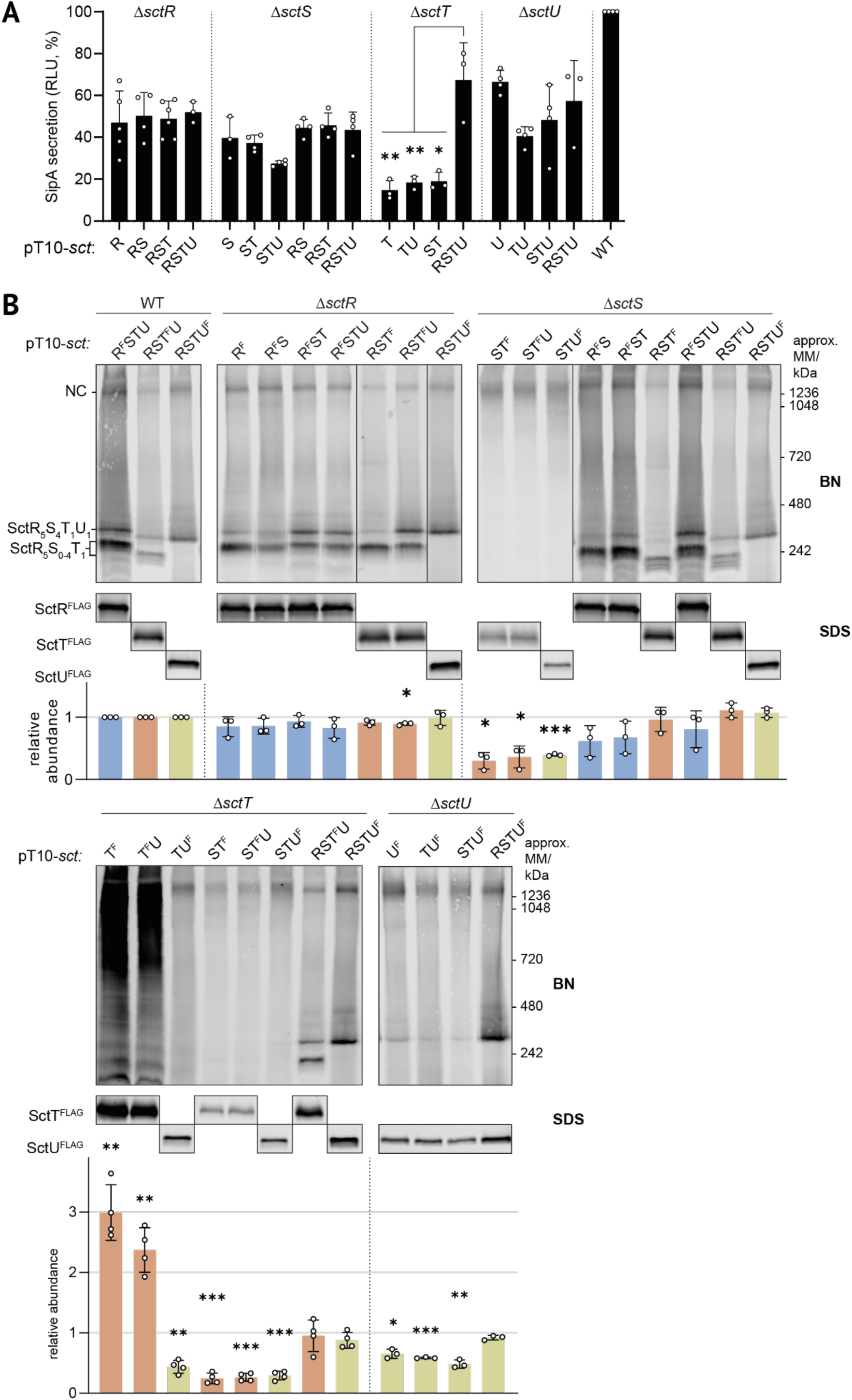
Plasmid-based complementation of chromosomal deletion mutants of individual export apparatus genes reveals a requirement for *sctT* expression within the conserved genetic organization of the export apparatus. **A.** SipA-NL secretion assay. Secretion function was assessed in *S*. Typhimurium strains with *sctR, sctS, sctT or sctU* deletions in a SipA-NL background after complementation with low copy number pT10 plasmid derivatives expressing export apparatus subunits. Luminescence values are scaled relative to a wild-type (WT) strain that expresses export apparatus proteins from the chromosome, which was set to a reference value of 100%. Data represent the mean of biological replicates (n ≥ 3), with standard deviations shown as error bars. Each complementation was compared with pT10-*sctRSTU* within each strain with an unpaired t-test. **B.** BN- and SDS-PAGE of solubilized crude membrane samples after plasmid complementation. SctR^FLAG^, SctT^FLAG^, and SctU^FLAG^ were visualized by immunoblotting of the triple FLAG tag, which is abbreviated as ‘F’. Bands corresponding to export apparatus assembly intermediates and the needle complex (NC) are annotated on the left of the BN-PAGE panels. The intensity of the SDS-PAGE bands was quantified using Image Studio software (Li-Cor, v5.2) and is shown as relative values compared to the samples of wild-type strain transformed with a pT10-*sctRSTU* plasmid containing the corresponding FLAG tags, which was set to a reference number of 1. Data of SctR^FLAG^, SctT^FLAG^, and SctU^FLAG^ are shown in blue, orange, and green, respectively. Data represent the mean of biological replicates (n ≥ 3), with standard deviations shown as error bars. P-values were calculated with one sample t-test, where the sample values were compared to the reference value of 1. ***, P ≤ 0.001, **, P ≤ 0.01, *, P ≤ 0.05. F, 3xFLAG.

We further investigated the relevance of the high synteny on the accumulation levels of export apparatus components by SDS-PAGE and on export apparatus assembly by blue native (BN)-PAGE (Zilkenat et al., 2024). SctR, SctT, and SctU, respectively, were analyzed using a 3xFLAG epitope tag, Western blotting and immunodetection as previously reported (Wagner et al., 2010). In the wild-type strains, all proteins were detected robustly by SDS-PAGE at their expected molecular mass (Fig. 2B). In the BN-PAGE, a distinct band of the respective proteins as part of the assembled needle complex was detected at above 1236 kDa. Subcomplexes of assembly intermediates containing SctRT (only weakly), SctRST and SctRSTU were detected around 242 kDa as previously reported (Fig. 2B) (Singh et al., 2021).

In the complementation of the Δ*sctR* mutant, expression of all constructs resulted in wild-type-like accumulation levels of the respective export apparatus components. Also the respective complexes occurred like in the wild-type strain (Fig. 2B).

Complementation of the Δ*sctS* mutant by plasmids lacking *sctR* resulted in SctT^FLAG^ and SctU^FLAG^ accumulation levels reduced to about 1/3rd of the wild-type, pointing at a severe stability defect of the respective proteins (Fig. 2B). Concomitantly, SctRST(U) subcomplexes were not found by BN-PAGE while protein complexes at the size of the full needle complex could be detected. Complementation of the Δ*sctS* mutant by plasmids encoding SctRS, SctRST and SctRSTU, respectively, yielded stable protein accumulation levels as well as the formation of all complexes. It is noteworthy that the intensity of the SctRSTU complex band was substantially stronger in the complementation with SctRST and even more with SctRSTU compared to complementation with SctRS only.

Complementation of the Δ*sctT* mutant by plasmids encoding SctT^FLAG^ or SctT^FLAG^U showed an up to three-fold overexpression of SctT^FLAG^ compared to the wild-type and a ladder of complex bands by BN-PAGE, presumably consisting of SctT multimers (Fig. 2B). Complementation of this mutant with every other combination except SctRSTU showed strongly reduced protein accumulation levels and no detectable formation of SctRST(U) subcomplexes.

Formation of the SctRSTU complex was also reduced in the Δ*sctU* mutant unless complemented with all four core export apparatus components (Fig. 2B).

Together, the results of the complementation analysis show that gene synteny is most critical for SctT and to a lesser extent also for SctS. Assembly of the SctRSTU subcomplex suffers if these proteins are not expressed in their genetic context. However, the surplus of proteins generated by plasmid-based complementation seems to yield a sufficient amount of functional needle complexes in most cases as a significant functional defect is limited to the complementation of the Δ*sctT* mutant. Expressing *sctT* without an upstream *sctS* gene results in SctT overexpression and futile multimerization of the protein.

### Scrambling of gene order and disconnecting *sctS* and *sctT* result in unproductive complex assembly

To further understand how the genetic organization affects export apparatus assembly, we generated constructs with scrambled orders of export apparatus genes. pT10 plasmids containing *sctSTRU*, *sctTRSU*, and *sctTSRU* complemented a Δ*sctRSTU S.* Typhimurium strain. Scrambling the gene order resulted in about 95% reduction in SipA-NL secretion compared to the wild-type gene order (Fig. 3A). A significant reduction in the accumulation levels of the respective proteins as well as inefficient NC and SctRSTU complex assembly were observed throughout with the exception of the overexpression of SctT and its concomitant multimerization in complementations with *sctT^FLAG^RSU* and *sctT^FLAG^SRU* (Fig. 3B).

**Figure 3.**
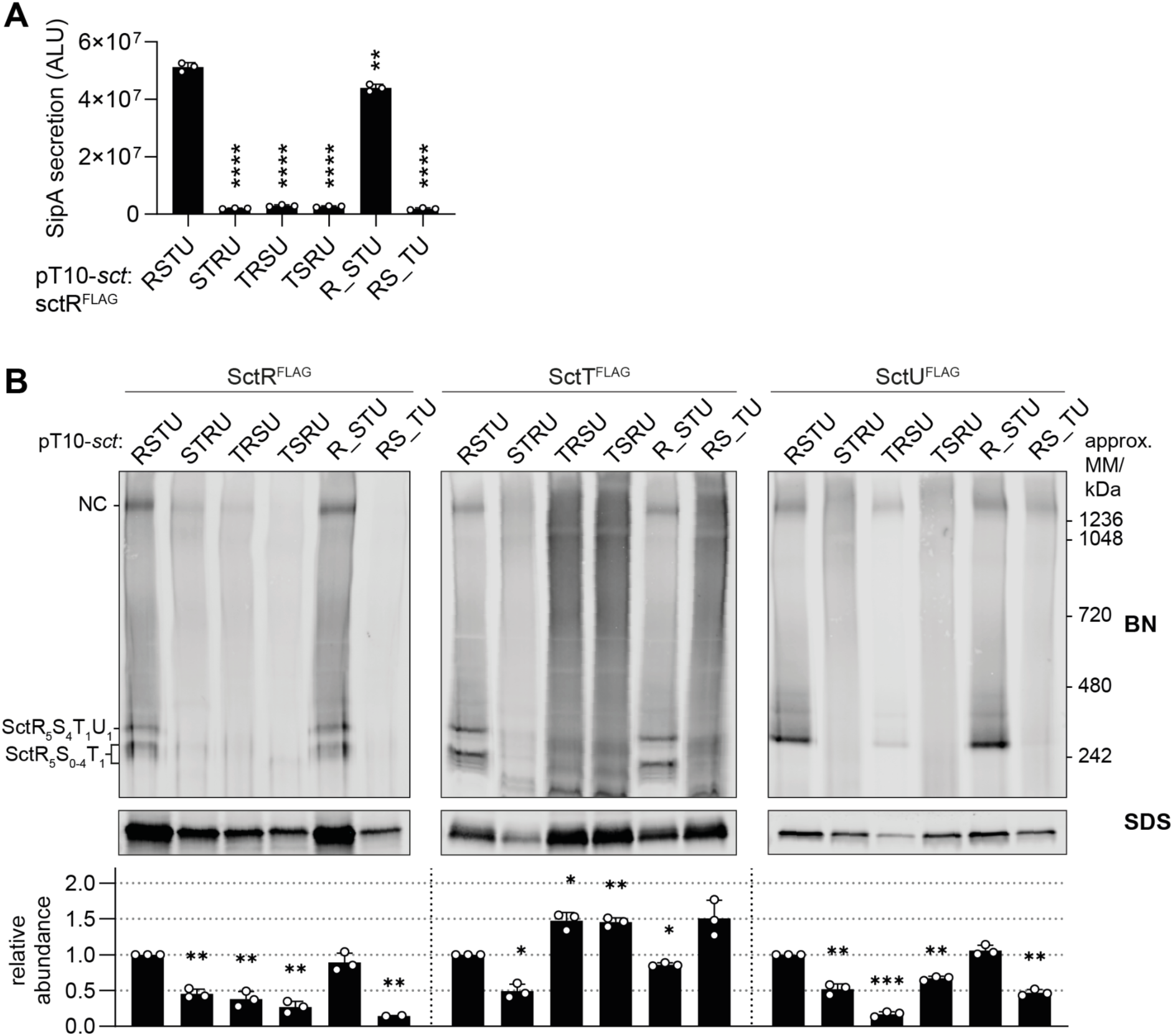
Disconnection of the *sctS*-*sctT* gene order impairs export apparatus assembly and type III secretion. **A.** SipA-NL secretion was measured in a *S*. Typhimurium Δ*sctRSTU* mutant strain with a SipA-NL background, which was transformed with pT10 plasmid derivatives with scrambled order (*sctSTRU, sctTRSU* and *sctTSRU*) or disconnection (*sctR_STU* and *sctRS_TU,* where 20 bp spacer is denoted as _) of export apparatus genes, each of which contained the 3xFLAG tag on *sctR*. P-values of each sample were calculated using an unpaired t-test, comparing them to the wild-type (*sctRSTU)* value. ALU, arbitrary luminescence unit. **B.** The same samples used in the secretion assay were analyzed by BN- and SDS-PAGE, which were performed as described in Fig. 2. The intensity of the SDS-PAGE bands was quantified using Image Studio software (Li-Cor, v5.2) and is shown as relative values to the wild-type gene order (*sctRSTU*) samples, which were set to a reference value of 1. The data represent the mean of biological replicates (n ≥ 2) with the standard deviations shown as error bars. P-values were calculated with one sample t-test, where the sample values were compared to the reference value of 1. ****, P ≤ 0.0001. ***, P ≤ 0.001, **, P ≤ 0.01.

A strict requirement for the conservation of gene order indicates that the connections between the genes may play a role in core export apparatus assembly. Towards testing this hypothesis, a 20 bp spacer (denoted as _) was introduced between SctR and SctT, and SctS and SctT, respectively. The respective native Shine-Dalgarno (SD) sequence of *sctS* or *sctT* was placed within the spacer at the native distance from the start codon. As SctU is not strictly required for NC assembly (Wagner et al., 2010) and *sctU* does not show high synteny relative to *sctRST* (Fig. 1B), the *sctT-sctU* gene connection was omitted from analysis. The spacer between *sctR* and *sctS* did not affect the assembly (Fig. 3B) and resulted only in a minor reduction (14%) of SipA-NL secretion (Fig. 3A). On the other hand, the spacer between *sctS* and *sctT* resulted in impaired assembly and secretion function (96% reduction) (Fig. 3A), as well as in SctT overexpression and aggregation, alongside a reduced accumulation of SctR and SctU (Fig. 3B). The effect of disconnecting *sctS* and *sctT* was even more pronounced when the spacer was introduced in the chromosome, with an increase in the SctT accumulation levels of 14-fold, highlighting the importance of the *sctS-sctT* gene connection (Supplementary Fig. 1). Together, these results indicate that the need for a particular connection between *sctS* and *sctT* is one critical aspect of the synteny of core export apparatus genes. Disruption of this connection leads to unregulated expression of SctT with concomitant multimerization, instability of the other core export apparatus components and a defect in the assembly of functional needle complexes.

### An mRNA stem-loop structure at the end of *sctS* inhibits independent translation of *sctT*

The observation of the overexpression of *sctT* in the absence of *sctS* immediately upstream points to a negative regulation of *sctT* expression by *sctS*. We hypothesized that this regulation could come from an inhibitory mRNA structure. Prediction of the mRNA structure (J. Singh et al., 2021) identified a stem-loop structure at the 3’ end of the *sctS* mRNA (Fig. 4A). As the SD sequence of *sctT* is located within the 3’ end of *sctS*, the predicted *sctS* stem-loop structure may restrict SD recognition by a ribosome and independent translation initiation of *sctT*.

**Figure 4.**
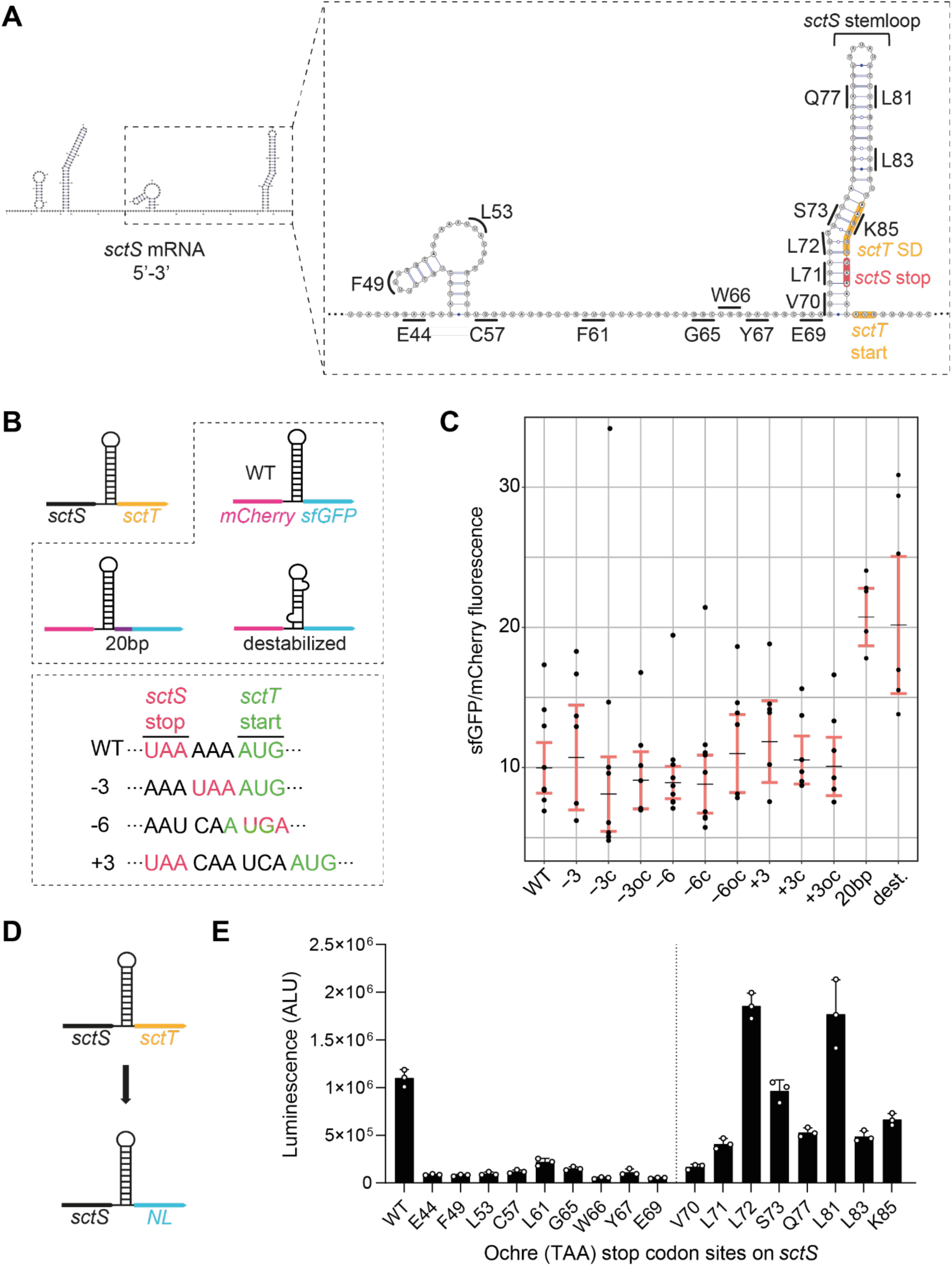
*sctS* and *sctT* are translationally coupled through a mRNA structure in *sctS*. **A.** Predicted secondary structure of the *sctS* mRNA (5’-3’) including the first 9 nucleotides of *sctT* was generated using SPOT-RNA2 (Singh et al., 2021). The Shine-Dalgarno (SD) sequence and start codon of *sctT* are indicated in orange, and *sctS* stop codon is indicated in red. Codons in *sctS* where the ochre stop codons were introduced are indicated on the graph. **B.** *sctS* and *sctT* genes were replaced with *mCherry* and *sfGFP* to use fluorescence as a readout for SctS and SctT expression, respectively. The stem-loop sequence of *sctS* was fused to *mCherry* (*mCherry:sctS*^S64-G86^), to mimic the native translational coupling between *sctS* and *mCherry*. Stem-loop destabilizing silent mutations (dest.; *sctS* L72 CTC::CTG, A82 GCG::GCA) or a 20 bp spacer (purple bar) were introduced to the pT12-*mCherry-sfGFP* construct as indicated in the top box. In the bottom box, mutations for varying the distance between the *sctS* stop and *sctT* start codons are shown, which were introduced to the pT12-*mCherry-sfGFP* construct. The distance between *sctS* stop and *sctT* start codons was reduced by 3 or 6 nucleotides (−3, -6) or increased by 3 (+3). Complementary ‘c’ mutations were further introduced on the other side of the stem-loop for maintaining the stem-loop structure. As a control, only the complementary mutation ‘oc’ was introduced without the distance mutations. **C.** Fluorescence of mCherry and sfGFP was acquired simultaneously, and the sfGFP/mCherry fluorescence ratio was calculated. In each plot, black lines mark the regression coefficients, and 95% confidence intervals are indicated in red. **D.** *sctT* was replaced with the NanoLuc (*NL*) gene to use luminescence as a readout for SctT expression. The pT12*-sctS-NL* plasmid was used as the basis for introducing the stop codons indicated in Fig. 4A. **E.** Luminescence measurements from the constructs with each ochre stop codon are shown in the bar graph. The vertical line separates codons located before and after the stem-loop. Three independent biological replicates were performed. The standard deviations are shown as error bars. ALU, arbitrary luminescence unit.

To investigate the role of the stem-loop in regulating *sctT* expression, we designed constructs in which *sctS* and *sctT* were replaced by *mCherry* and *sfGFP*, respectively, and were the *sctS* stem-loop sequence was fused to *mCherry* to mimic the native gene connection between *sctS* and *sctT* (Fig. 4B). The use of fluorescent proteins ensures that results are not influenced by membrane targeting and insertion, the stability of individual proteins, or complex assembly. We then introduced silent mutations to destabilize the stem-loop, and the aforementioned 20 bp spacer between the *mCherry* and *sfGFP* genes as described above, recapitulating the disconnection of *sctS* and *sctT* genes. Analysis of the fluorescence ratio of mCherry and sfGFP from both stem-loop destabilization and spacer mutants revealed a significant increase in sfGFP fluorescence compared to mCherry (Fig. 4C). This result indicates that the *sctT* SD sequence is concealed by the stem-loop structure and may require melting by a ribosome that translates SctS. This mechanism would couple the two genes translationally.

To test how far a ribosome needs to melt this stem-loop structure to translate *sctS*, we introduced ochre stop codons along the sequence of *sctS* (Fig.4A) and measured the luminescence output of NanoLuc luciferase (NL) that replaced the *sctT* gene (Fig. 4D). Introducing stop codons upstream of the stem-loop resulted in a strongly diminished luminescence compared to the wild-type. On the other hand, introducing stop codons within the stem-loop led to NL expression. Starting from position L72, which base-pairs with the *sctT* SD sequence, a substantial increase in the NL expression was observed upon stop codon introduction (Fig. 4E). These results indicate the presence of a strict translational coupling between *sctS* and *sctT: sctT* is only translated when the ribosome translating *sctS* melts the stem-loop structure and allows access to the *sctT* SD. Otherwise, the *sctS* stem-loop inhibits *sctT* SD recognition by a ribosome and independent *sctT* translation.

Previous studies on translational coupling have shown that the distance between the stop codon of the upstream gene and the start codon of the downstream gene affects translational coupling efficiency (van de Guchte et al., 1991). To investigate if this distance is fine-tuned for the translational coupling of *sctS* and *sctT*, we made use once more of the *mCherry-sctS_stem-loop_-sfGFP* plasmid (Fig. 4B). The distance between *sctS* stop and *sctT* start codons was either increased or decreased by 3 nt (denoted as +3 and -3). Additionally, overlapping stop and start codons (AUGA) were generated (denoted as -6). In order to maintain the stem-loop structure upon changing the distance, complementary mutations (denoted as ‘c’) were introduced to the opposite strand of the stem-loop. The sole effect of the complementary mutations was tested as a control (denoted as ‘oc’). The ratio between mCherry and sfGFP fluorescence did not change upon varying the distance between *mCherry* stop and *sfGFP* start codons (Fig. 4C). Together, these results suggest that the distance between the *sctS* stop and *sctT* start codons is not optimized towards a particular translational coupling, eventually leading to the required 4:1 stoichiometry of the two complex components.

### Regulation of *sctT* translation by the stem-loop structure is independent of the gene in front of *sctT*

Proper folding and assembly of protein complexes often depend on co-translational processes (Shiber et al., 2018). Since export apparatus subunits require each other for stability (Wagner et al., 2010), co-translation may help circumvent an inherently unstable monomeric state, allowing the complex to form more efficiently.

As SctS and SctT proteins interact and are translationally coupled, we wondered if SctS and SctT require each other for co-translational assembly. To this end, *mCherry* was inserted upstream of *sctT*, coupling *sctT* and *mCherry* in a manner analogous to the native *sctS-sctT* connection and at the same time separating *sctS* from *sctT*. The *sctS* stem-loop sequence was fused to the 3’ end of the *mCherry* gene either in the presence or absence of *sctS* (Fig. 5A).

**Figure 5.**
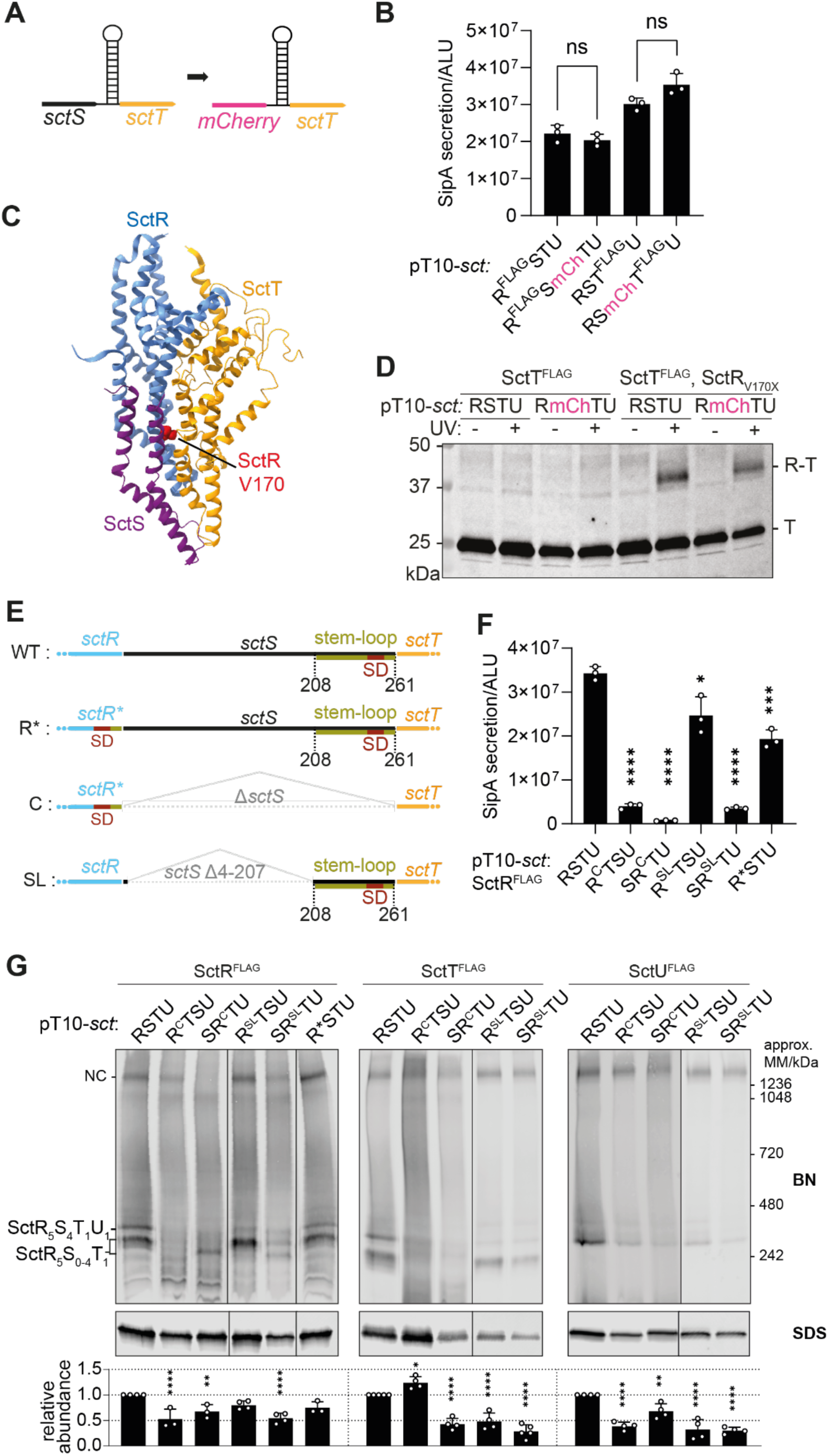
Translational coupling through the *sctS* stem-loop and ordered gene organization prevents SctT overexpression and ensure sequential assembly of the export apparatus. **A.** Schematic diagram showing the translational coupling between *mCherry* and *sctT*, established by fusing a remnant of the *sctS* stem-loop sequence to *mCherry* (*mCherry:sctS*S64-G86). **B.** SipA-NL secretion was measured in the *S*. Typhimurium Δ*sctRSTU* mutant strain with a SipA-NL background, which was transformed with pT10 plasmid derivatives harboring either the wild-type gene order or the *sctRS-mCh-sctTU* construct, with the *mCherry-sctT* connection as described in Fig. 5A. P-values of each sample were calculated using an unpaired t-test. Three independent biological replicates were performed. The standard deviations are shown as error bars. ALU, arbitrary luminescence unit. ns, not significant.**C.** Partial structure of the export apparatus (PDB 6F2D) (Kuhlen et al., 2018) showing the SctR-SctS-SctT interface, with the SctR V170 residue highlighted in red. SctR, SctS, and SctT are shown in blue, purple, and orange, respectively. **D.** *In vivo* photo-crosslinking of SctR and SctT in *S*. Typhimurium Δ*sctRSTU* mutant strain, transformed with pT10 plasmid derivatives harboring either the wild-type gene order or *sctRS-mCh-sctTU* construct, with and without amber stop codon sites (indicated as X). **E.** Schematic diagram showing constructs with different *sctR-sctT* arrangements including the wild-type (WT), ‘directly <u>c</u>onnected’ (C), and ‘stem-loop’ (SL). In the wild-type, the *sctS* stem-loop (olive green) contains the *sctT* Shine-Dalgarno (SD) sequence (red). In the C connection, *sctS* was deleted and the SD was placed in *sctR*. In SL connection, *sctS*4-207 was deleted, but the stem-loop sequence was retained. R* mutation (SctR A223K and T224G) were introduced to the C connection to accommodate *sctT*’s SD within *sctR*. R* alone was included as a control for the C connection. **F.** SipA-NL secretion was measured in *S*. Typhimurium *ΔsctRSTU* mutant strain with a SipA-NL background, which was transformed with pT10 plasmid derivatives harboring either the wild-type gene order or C and SL constructs in *sctRTSU* and *sctSRTU* gene orders as described in Fig. 5E. The statistical analysis was conducted as described in Fig. 5B. **G.** BN and SDS-PAGE were performed as described in Fig. 2. The intensity of the SDS-PAGE bands was quantified using Image Studio software (Li-Cor, v5.2) and is shown as relative values compared to the wild-type gene order (*sctRSTU*) samples, which were set as 1. The data represent the mean of three biological replicates, with the standard deviations shown as error bars. P-values were calculated using one way ANOVA. ****, P ≤ 0.0001. ***, P ≤ 0.001, **, P ≤ 0.01.

Then, we assessed the SipA-NL secretion after complementation of the Δ*sctRSTU* mutant with the pT10-*sctRSmcherryTU* construct. Surprisingly, translational coupling by *mCherry-sctT* instead of *sctS-sctT* did not impact the secretion function, indicating that co-translational interaction between SctS and SctT is not the critical point of export apparatus assembly and the reason for translational coupling of these two proteins (Fig. 5B).

In addition, we assessed the assembly of the SctR_5_T_1_ subcomplex with *mCherry-sctT* translational coupling and without SctS via *in vivo* photo-crosslinking. This method relies on the genetic encoding of the artificial UV-reactive amino acid *para*-benzoyl-phenylalanine (*p*Bpa) by suppressed amber stop codons (UAG) that can be introduced by site-directed mutagenesis at any protein position of interest (Farrell et al., 2005; N. Singh et al., 2021). Upon UV irradiation, *p*Bpa forms free radicals and crosslinks to closely interacting proteins. The observation of a *p*Bpa crosslink, e.g. by Western blotting of crosslinked proteins, ascertains that a specific protein-protein interaction has occurred. To assess SctR-SctT assembly, we replaced V170 of SctR by pBpa; SctR_V170X_ was reported previously to crosslink to SctT (Fig. 5C) (Dietsche et al., 2016). The absence of SctS did not affect the crosslinking efficiency of SctR_V170X_ to SctT (Fig. 5D), supporting the notion that co-translational interaction of SctS and SctT is not relevant for assembly of the core export apparatus as long as SctT expression is regulated by translational coupling via the *sctS* mRNA stem-loop.

While the assembly of the SctR_5_T_1_ subcomplex proves to be independent of the SctS protein, SctS requires SctR_5_T_1_ for its assembly. In order to evaluate how critical translational coupling of *sctS* and *sctT* is for SctS assembly onto SctR_5_T_1_ complexes, we designed constructs in which *sctS* was placed upstream or downstream of the *sctRT* genes (*sctSRTU* or *sctRTSU*, respectively). In the designs of the *sctRT* genes, we either maintained *sctT* translational coupling through a remnant *sctS* stem-loop (denoted as SL) or connected the *sctR* and *sctT* genes just by the original TAA AAA ATG sequence without the regulating stem-loop but providing the *sctT* SD within the *sctR* gene (denoted as C for directly Connected) (Fig. 5E). The A223K and T224G mutations of SctR to accommodate *sctT’s* SD within *sctR* (denoted R*STU in Figs. 5E-G) did not have a substantial impact on protein accumulation levels, complex assembly and type III secretion function. However, SipA-NL secretion was strongly reduced when *sctS* was upstream of *sctRT* (*sctSRTU*) in both C and SL connections (Fig. 5F). When expressing SctS behind SctRT, only the construct that included the *sctS* stem-loop between *sctR* and *sctT* (*sctR^SL^TSU*) successfully complemented type III secretion function while the direct connection of sctR and sctT failed to do so (Fig. 5F). Together, this suggests that the direct availability of the SctR_5_T_1_ subcomplex is important for the assembly of SctS.

In BN-PAGE, the constructs expressing the directly connected *sctR-sctT* genes without a *sctT*-controlling stem-loop produced a strongly altered banding pattern of subcomplexes with much reduced amounts of assembled needle complexes (Fig. 5G). A mild SctT overexpression and multimerization phenotype was observed when expressing *sctR^C^TSU*, while, with the exception of SctR in *sctR^SL^TSU*, all tested export apparatus components showed reduced accumulation levels in the different mutants pointing at compromised assembly and stability. Concomitantly, only expression of the *sctR^SL^TSU* construct resulted in wild type-like band intensities of SctR^FLAG^-containing subcomplexes.

In summary, our results reveal two important aspects in the core export apparatus assembly: First, *sctT* requires translational coupling through the stem-loop for appropriate expression levels, but the nature of the coupled gene in front is not important. Second, it seems to be critical for *sctS* to be translated after *sctR* in order to form SctRST complexes efficiently.

### Reducing *sctT* translation with inefficient start codons alleviates effect of uncoupled translation

Since translational coupling through the stem-loop prevented SctT overexpression and multimerization, we investigated whether the requirement for translational coupling can be waived by reducing *sctT* translation levels. It is known that translation levels correlate with the start codon usage (O’Donnell & Janssen, 2001), and that the alternative start codons GTG and TTG are used in 14% and 3% of *E. coli* genes, respectively, whereas ATG is used in 82% (Blattner et al., 1997; Hecht et al., 2017). Therefore, in the constructs that led to SctT overexpression, namely, *sctTRSU* and *sctTSRU*, we replaced the ATG start codon of *sctT* with two alternative start codons, GTG and TTG. The use of the TTG start codon significantly reduced SctT levels compared to the ATG start codon (Fig. 6A). In the BN-PAGE, high and low molecular weight SctT-containing complexes were reduced when the alternative start codons were used, with the TTG start codon having a more pronounced effect than GTG, reflecting their translation efficiency (Fig. 6A). Having SctT overexpression reduced also alleviated the secretion defect by 2-fold with TTG and up to 4-fold with GTG compared to the ATG start codon (Fig. 6B). In the case of the *sctTSRU* construct, having a GTG start codon restored the secretion function up to 82% of the wild-type gene order, compared to18% with the ATG start codon (Fig. 6B). We also observed that SctT overexpression caused growth attenuation of the expressing bacteria (Fig. 6C). Reducing SctT expression with alternative start codons restored growth to wild-type levels (Fig. 6C). Taken together, these results suggest that reducing SctT overexpression can compensate for the lack of translational coupling between *sctS* and *sctT*, and the associated problems in assembly, secretion and growth.

**Figure 6.**
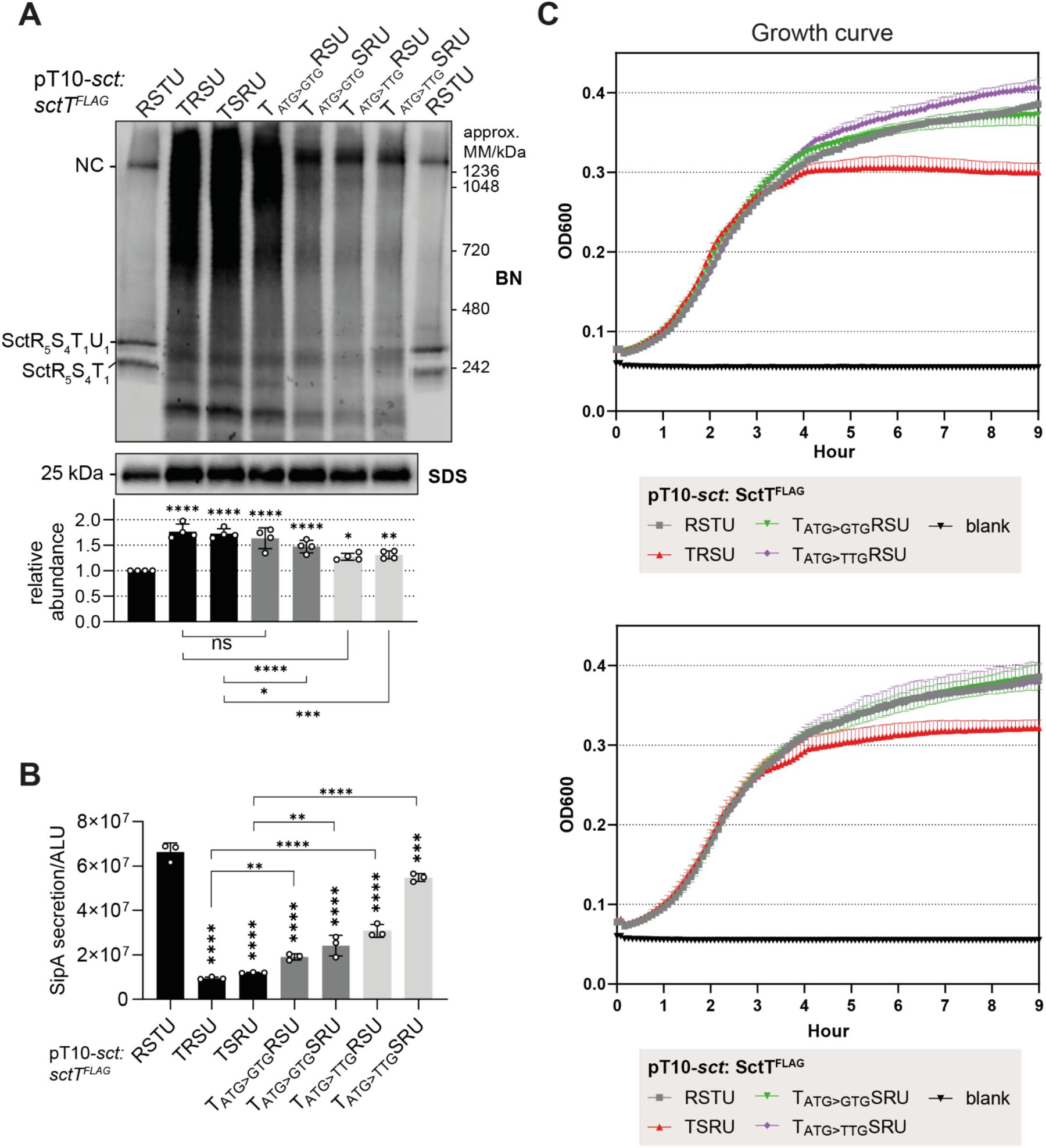
Reducing the translation of SctT compensates for the lack of translational coupling regulation through *sctS* stem-loop. **A.** *S.* Typhimurium Δ*sctRSTU* mutant strain in SipA-NL background was transformed with pT10 plasmids *sctTRSU* or *sctTSRU,* with ATG, GTG or TTG start codons on *sctT.* BN- and SDS-PAGE were performed as described in Fig. 2. The intensity of the SDS-PAGE bands was quantified using Image Studio software (Li-Cor, v5.2) and are shown as relative values to the wild-type gene order (*sctRSTU*) samples, which were set as a reference value of 1. The data represents the mean of four biological replicates with the standard deviations shown as error bars. P-values of each sample were calculated using one way ANOVA. P-values calculated from comparisons against the wild-type (*sctRSTU)* are shown above each bar graph. The comparisons between the mutants are shown below the graph with a line with the P-value. **B.** SipA-NL secretion of the *S.* Typhimurium transformants. The data represents the mean of ≥3 biological replicates with the standard deviations shown as error bars. P-values calculated from comparisons against the wild-type (*sctRSTU)* are shown above each bar graph. The comparisons between the mutants are shown with a line with the P-value. ALU, arbitrary luminescence unit. **C.** The growth curve of *S.* Typhimurium transformants was measured at OD600 absorbance every 5 minutes for 9 hours. The data represents the mean of three biological replicates with the standard deviations shown as error bars. ****, P ≤ 0.0001. ***, P ≤ 0.001, **, P ≤ 0.01, *, P ≤ 0.05.

### Strict regulation of *sctT* translation by the mRNA stem-loop prevents futile polymerization of SctT

Why is SctT overexpression so detrimental to export apparatus assembly and even results in affecting bacterial growth? To address this, we had a closer look at the structure of the export apparatus. SctT resembles a unit of SctR and SctS, which makes the export apparatus a pseudo-hexameric pore (Kuhlen et al., 2018). Hence, it is well conceivable that SctT assembles into SctT-only multimers in the absence of stem-loop regulation (Fig. 7A). To validate self-assembly of SctT, we performed *in viv*o photo-crosslinking of SctT with a pBpa mutant (SctT_F132X_) that reveals SctT-SctS crosslinks in the wild-type (Dietsche et al., 2016) (Fig. 7AB). When SctT was overexpressed due to the lack of control by the stem-loop, these crosslinks were absent. Instead, SctT was crosslinked with SctT, forming homo-oligomers (Fig. 7B). This result supports that the ladder observed by BN-PAGE upon SctT overexpression corresponds to SctT multimers (Fig. 6A). This self-assembly likely results in the formation of ungated leaky pores that impair membrane integrity and consequently bacterial growth. Alternative to the assumed helical self-assembly of SctT, also a planar penta- or hexameric pore formation is conceivable as predicted by AlphaFold multimer (Supplementary Fig. 2). Due to the propensity of SctT for futile self-assembly, a tight control by the stem-loop may become crucial to ensure that only a few copies of SctT are translated, which then are swiftly assembled onto SctR_5_ to form SctR_5_T_1_.

**Figure 7.**
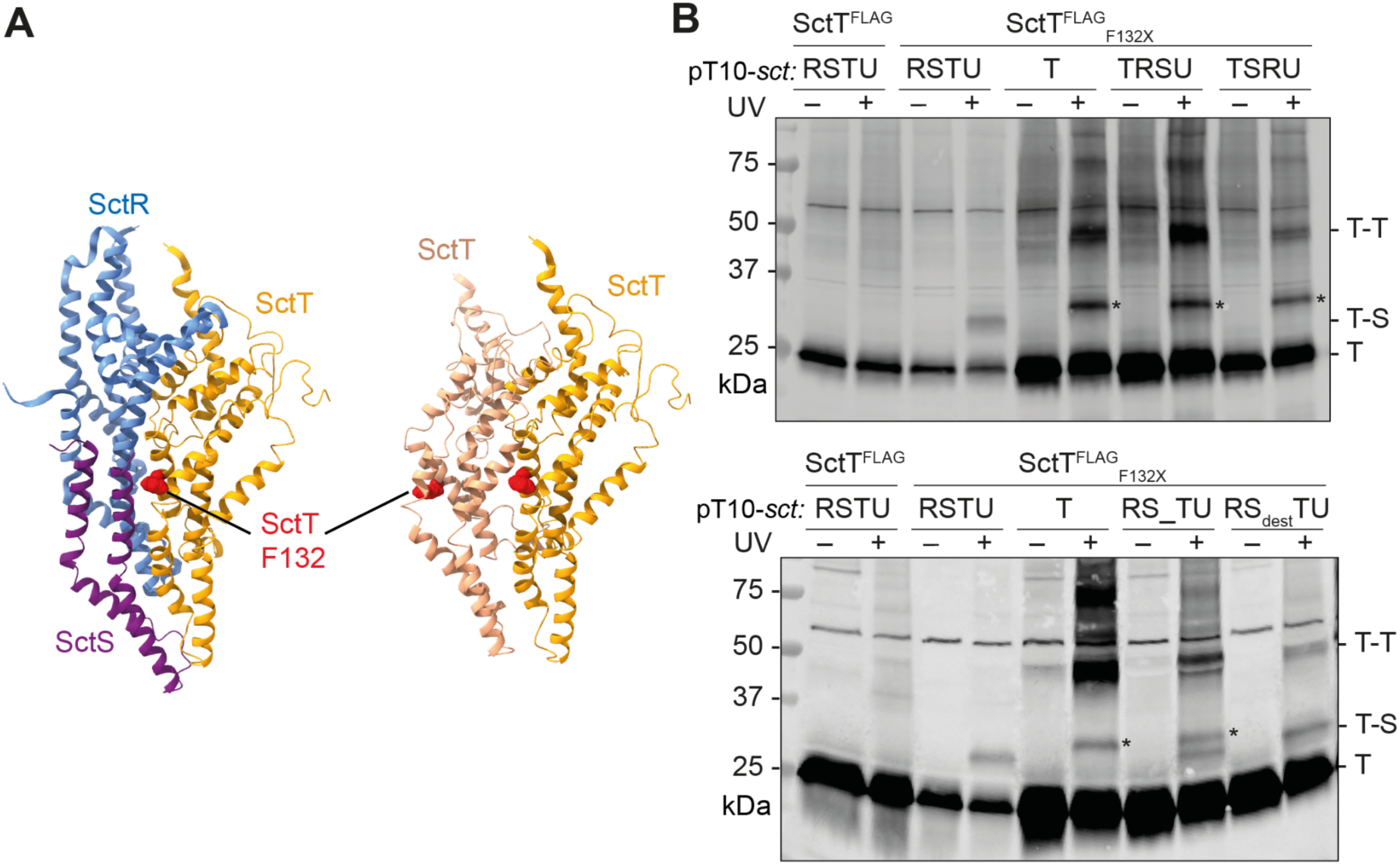
Uncontrolled expression results in futile SctT multimerization. **A.** Partial structure of the export apparatus (PDB 6F2D) (Kuhlen et al., 2018), showing the SctR-SctS-SctT interface (left). SctR, SctS, and SctT are shown in blue, purple, and orange, respectively. A hypothetical SctT-SctT interface is shown on the right, with SctT molecules in shades of orange. SctT F132 is indicated in red. **B.** Futile SctT-SctT interactions upon SctT overexpression as shown by *in vivo* photo-crosslinking. *S.* Typhimurium Δ*sctRSTU* mutant strains expressing pT10 plasmid derivatives with or without crosslinking residue (SctT F132X, where X denotes an amber stop codon encoding pBpa) were irradiated or not with UV irradiation for 30 min. Crude membrane fractions were subjected to SDS-PAGE and analyzed by Western blotting detecting 3xFLAG tag. Cross-linked interacting partners of SctT are indicated to the right of each blot. Asterisk (*) indicates a SctT-specific band, which may represent a crosslink of a SctT degradation product.

### Conservation of *sctS* stem-loop structure in other T3SS

Our study has identified translational coupling of *sctST* as a crucial regulatory mechanism for export apparatus assembly in SPI-1 T3SS in *S.* Typhimurium. Given the conserved synteny in the genes encoding the export apparatus subunits among pathogens with T3SS (Fig. 1A and 1C), we hypothesized that control of *sctT* translation through a *sctS* mRNA stem-loop may be a more universal mechanism to facilitate faithful assembly of the core export apparatus. Indeed, we found similar *sctS* mRNA stem-loop structures that are predicted to be present around *sctT* SD sequences of other T3SS (Supplementary Fig. 3), supporting our notion. As the *sctS* mRNA structures are machine learning-based predictions, the presence of *sctS* stem-loop structures and their regulatory effects on *sctT* translation and export apparatus assembly awaits further experimental validation for other pathogens.

## Discussion

The T3SS is an intricate protein complex machinery that requires highly coordinated assembly of its subunits and subcomplexes, which involves a variety of regulatory measures. In this study, we investigated the assembly regulation of the export apparatus. Our findings reveal that high synteny in export apparatus genes relates to successful export apparatus assembly. A key regulatory motif in the conserved genetic organization of the export apparatus is the translational coupling between the *sctS* and *sctT* genes. This translational coupling occurs through an mRNA stem-loop structure, which conceals the Shine-Dalgarno (SD) sequence of the *sctT* gene. The stem-loop inhibits *de novo* translation of SctT by a new ribosome, allowing SD recognition only through translational coupling with *sctS*. In absence of translational coupling via the stem-loop, SctT is overexpressed and assembles into futile multimers (Fig. 8).

**Figure 8.**
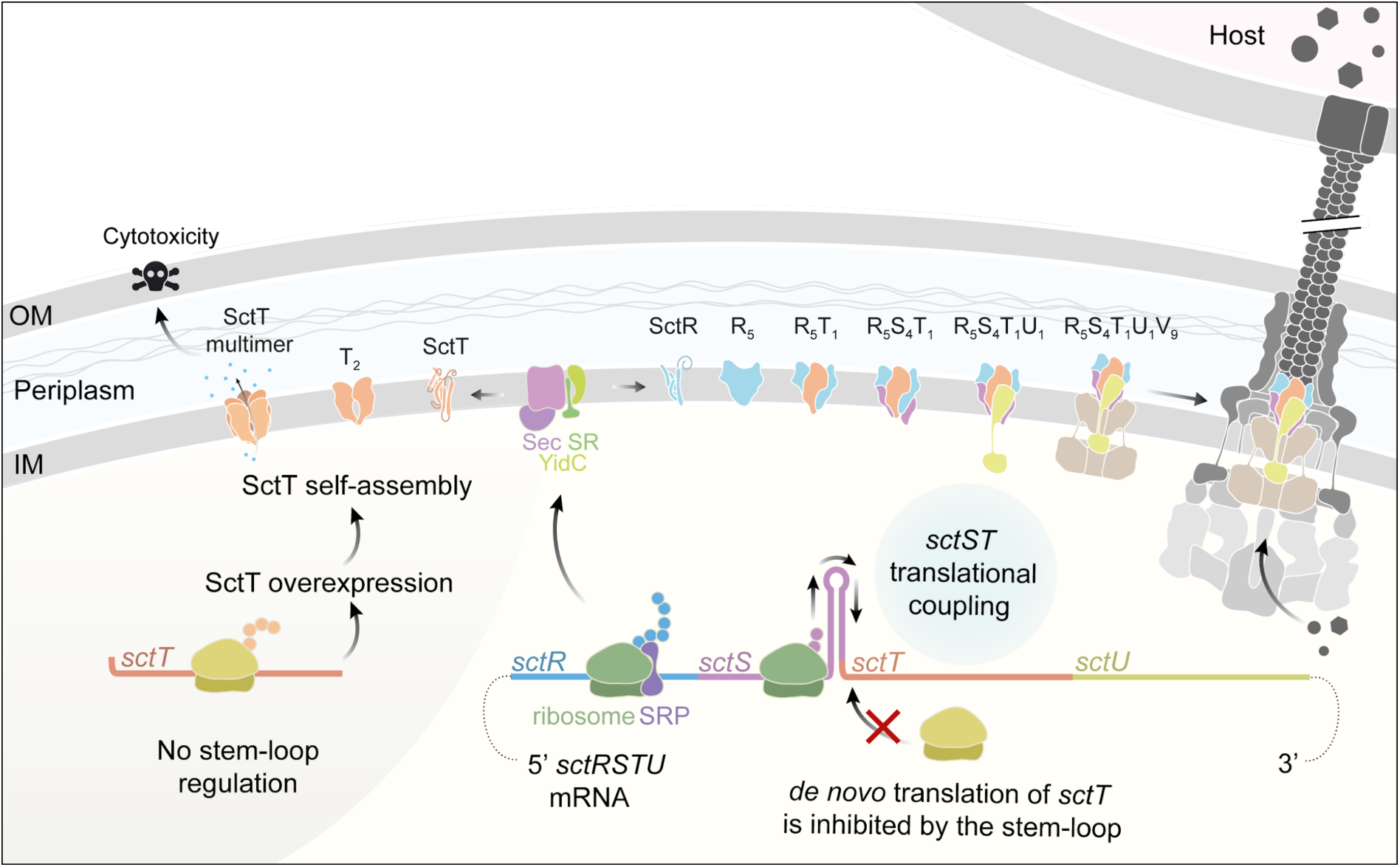
Summary of export apparatus assembly and *sctS*-*sctT* translational coupling as a fine-tuning mechanism. The export apparatus genes *sctRSTU* are transcribed in one transcript, which exhibits a high degree of synteny. The expression of *sctT* is under tight control involving translational coupling with *sctS* through the *sctS* mRNA stem-loop structure. Recognition of *sctT* SD by a ribosome is inhibited unless the ribosome translating *sctS* melts the stem-loop. In the absence of control by the stem-loop, SctT is overexpressed and self-assembles into futile multimers. SctT self-assembly under conditions of unregulated SctT expression disrupts the export apparatus assembly and causes cytotoxicity. Control of *sctT* translation by the *sctS* stem-loop ensures appropriate levels of SctT and swift complexation of SctT with the SctR pentamer. The SctR5T1 subcomplex subsequently recruits SctS and SctU subunits for finalizing the helical assembly of the export apparatus. Successfully assembled export apparatus then nucleates the further assembly of the needle complex base. SRP: signal recognition particle. SR: SRP receptor. OM, outer membrane. IM, inner membrane. The injectisome illustration was adapted from (Pais et al., 2023).

Our results stand in contrast to a study reporting that complete recoding of the SPI-1 operon, eliminating all internal regulatory elements for assembly, still permitted formation of needle complexes (Song et al., 2017). However, this study only provides a qualitative assessment of complex formation with no quantitative information on the efficiency of T3SS assembly from the synthetic T3SS operon, in which all components were overexpressed and thus available in surplus. In physiological context, cellular resources are limited, and the T3SS, a massive and energy-costly complex, requires its components to be produced in strict stoichiometric ratios. Supporting this, ribosome profiling in *E. coli* has shown that subunits of protein complexes are typically synthesized in precise proportion to their final stoichiometric ratios (Li et al., 2014). In line with this, our results clearly demonstrate that the conserved gene order *sctRSTU* is important for stoichiometric expression of the export apparatus subunits. This is especially important for SctT, whose overexpression is detrimental to pathogen fitness, and it ultimately ensures efficient and robust assembly of the export apparatus.

The propensity of SctT to self-interact can be inferred from the structure of the export apparatus. SctT alone resembles a structural combination of SctR and SctS and constitutes a unit within the pseudo-hexameric SctR_5_S_4_T_1_ pore (Kuhlen et al., 2018). It is thus conceivable that SctT forms pore-like assemblies similar to the export apparatus. However, in the absence of the gating mechanism in the export apparatus (Hüsing et al., 2021; Kuhlen et al., 2018; Miletic et al., 2021), such aberrant SctT pores may lead to membrane leakage and growth defects, as observed under SctT overexpression conditions (Fig. 6). Given that the export apparatus is a highly conserved component of the T3SS and shares high structural similarity with its homologs across closely related bacterial species as well as in the flagellar system (Johnson et al., 2019; Lunelli et al., 2020), it is plausible that SctT homologs likewise possess a propensity for self-assembly and hence require regulations for proper complex formation.

The fate of SctS molecules during formation of SctR_5_T_1_ subcomplexes remains an intriguing question. A previous study by our lab demonstrated that SctT can stably assemble into the export apparatus only when salt bridges are established within the SctR_5_T_1_ subcomplex (N. Singh et al., 2021). Since SctS is translated before SctT, it is conceivable that SctS molecules diffuse away from the site of export apparatus assembly. A gene order matching the assembly order would therefore seem more logical unless there is another reason for *sctS* to be positioned upstream of *sctT*. Studies on translational coupling have shown that proteins from downstream genes are often chaperoned by proteins from upstream genes, aiding in correct folding (Basu et al., 2004; Nakatogawa et al., 2004; Praszkier & Pittard, 2002). However, our crosslinking data showed that SctS is dispensable for SctR_5_T_1_ subcomplex formation, arguing against the possibility that SctS functions as an intramembrane chaperon for SctT (Fig. 5D). On the other hand, we showed that expressing the export apparatus genes in the assembly order, with wild-type level SctT expression, still fails to recapitulate the robustness of the wild-type gene order (Fig. 6A). This indicates that the conserved gene order indeed fine-tunes the assembly and suggests a possible additional role of SctS in this process. It is possible that SctS transiently interact with the preformed SctR pentamer and stabilize its assembly by forming salt bridges with SctT. This hypothesis requires further investigation.

Given this inherent risk of SctT’s self-assembly, it is conceivable that maintaining the *sctS*-*sctT* synteny serves to preserve the *sctS* stem-loop, which in turn tightly regulates SctT translation and ensures proper assembly of the export apparatus. While our study shows that the *sctS* stem-loop regulates SctT translation via translational coupling in *Salmonella*, it may well be that the *sctT* stem-loop has different functions and mechanisms of action in other bacteria. For instance, in *Yersinia*, the presence of an *sctS* stem-loop has been reported, which harbors the SD sequence of *sctT,* like in *Salmonella* (Pienkoß et al., 2022). However, the *Yersinia sctS* stem-loop acts as an RNA thermometer, enabling translation of SctT upon reaching the host temperature (Pienkoß et al., 2022). An RNA thermometer is a regulatory motif frequently observed in T3SS regulation in *Yersinia,* as it aligns with *Yersinia’s* lifestyle and infection process (Hoe & Goguen, 1993; Pienkoß et al., 2021, 2022).

Together, we conclude that the synteny of the export apparatus genes provides a critical regulation for helical assembly of the pseudo-hexameric complex. Due to its unique structural symmetry, the export apparatus subunit SctT is prone to self-assembly and the *sctS* stem-loop provides a strict expression control via translational coupling between *sctS* and *sctT*.

## Materials and methods

### Materials

Most chemicals were purchased from Sigma-Aldrich, Novagen, New England Biolabs, Merck, Carl Roth, Becton Dickinson, or Applichem, unless otherwise specified. *Para*-benzophenylalanine (*p*Bpa) was purchased from Bachem. Lauryl maltose neopentyl glycol (LMNG) was from Anatrace Products LLC. SERVAGel™ TG PRiME™ 8-16% precast gels were purchased from SERVA. The monoclonal M2 anti-FLAG primary antibody was purchased from Sigma-Aldrich. The goat anti-mouse IgG secondary antibody DyLight^TM^ 800 conjugate was from Thermo-Fisher (SA5-35521). Plasmids and primers used in this study are listed in Supplementary Table 1. Primers were synthesized by Eurofins Genomics.

### Bacterial strains, plasmids and growth condition

Bacterial strains and plasmids used in this study are listed in Supplementary Table 1. Unless otherwise specified, bacteria were grown at 37°C in liquid medium shaken at 180 rpm under low aeration. Cultures were supplemented with 1 mM arabinose and 1 mM rhamnose to induce expression of pBAD24-hilA and pT10 plasmid derivatives, respectively (Wagner et al., 2010).

### I*n silico* analysis of export apparatus synteny

The Cblaster program was used to identify homologs of export apparatus subunits and to visualize the genetic organization of the corresponding genes (Gilchrist et al., 2021). The search module was run with the following parameters: a minimum identity of 15.0%, minimum coverage of 25.0%, maximum E-value of the BLAST hit of 0.01, and a maximum distance of 20,000 bp between any two hits in a cluster. The BLAST search was performed against the NCBI *refseq_protein* database using the following query sequences from *S*. Typhimurium strain LT2: NP_461808.1, NP_461809.1, NP_461810.1, and NP_461811.1. Search results were filtered using the list of reference bacterial genomes from NCBI, and the resulting data were further processed to visualize the distribution of genetic organization patterns. Taxonomic information of each hit was retrieved from NCBI Taxonomy database and visualized as a taxonomic tree using Interactive Tree of Life (iTOL) v7.2.1 (Letunic & Bork, 2024). A comprehensive list of BLAST hits, including BLAST scores, genome annotations, and taxonomic assignments, is provided in Supplementary Table 2.

### Secretion assay

Secretion assay was performed as previously described by Westerhausen et al. (Westerhausen et al., 2020). *S.* Typhimurium strains with the SipA-NL background were grown at 37°C for 5 h in LB medium supplemented with 0.3 M NaCl and appropriate antibiotics for plasmid selection under low aeration. Culture supernatants were collected by centrifugation at 10,000 × *g* for 2 min. Nano-Glo® luciferase substrate and Nano-Glo® luciferase assay buffer (Promega) were mixed at 1:50 ratio and then combined 1:1 with the culture supernatant. After a brief incubation, luminescence was measured using a microplate reader (Tecan Spark).

### *In vivo* photocrosslinking

*S.* Typhimurium strains were grown at 37°C for 5 h in LB medium supplemented with 0.3 M NaCl and appropriate antibiotics for plasmid selection under low aeration. Cultures were supplemented with the artificial amino acid para-benzoyl phenyl alanine (*p*Bpa) to a final concentration of 1 mM. 2 OD units of bacterial cells were harvested by centrifugation at 4,000 × *g* for 5 min and washed once with 1 mL 1x PBS. Cells were resuspended in 1 mL 1x PBS and transferred into 6-well cell culture dishes. UV irradiation (λ = 365 nm) were performed for 30 min at 4°C on a UV transilluminator table. Cells were then pelleted by centrifugation for 2 min at 10,000 × *g* and 4°C. The cell pellets were either immediately subjected to SDS-PAGE or stored at -20°C until further use.

### SDS-PAGE and Immunoblotting

For protein detection, samples were subjected to SDS-PAGE using SERVAGel™ TG PRiME™ 8–16% 12-sample-well precast gels (SERVA Electrophoresis) and transferred onto an Immun-Blot® polyvinylidene difluoride (PVDF) membrane (0.2 µm, Bio-Rad). Membranes were blocked with 1x Blue Block PF (SERVA electrophoresis), then probed with Monoclonal M2 anti-FLAG antibody (1:5,000 in TBS-T, Sigma-Aldrich), followed by goat anti-mouse IgG DyLight 800 conjugate (1:10,000 in TBS-T, Thermo Pierce). Scanning and image analysis was performed with a Li-Cor Odyssey scanner and Image Studio software (Li-Cor, v5.2).

### Crude membrane preparation

Crude membranes were prepared as reported previously (Zilkenat et al., 2024). 8 OD units of *S*. Typhimurium cultures were resuspended in 750 μl lysis buffer consisting of buffer K (50 mM triethanolamine, 250 mM sucrose, 1 mM EDTA, pH 7.5) supplemented with 1 mM MgCl_2_, 10 μg/ml DNAse I, 2 mg/ml lysozyme, 1:100 protease inhibitor cocktail (P8849, Sigma-Aldrich) and 2 mM EDTA. After incubation at 4°C for 30 min, samples were mixed with glass beads (150-212 µm) and lysed by bead milling for 2 minutes (SpeedMill Plus, Jena Analytics). Cell lysates were centrifuged at 1,000 × *g* for 1 min at 4°C, and the supernatant was collected. The glass beads were resuspended with 1 ml buffer K, centrifuged at 1,000 × *g* for 1 min at 4°C, and the supernatant was pooled. Cellular debris were removed by centrifugation at 10,000 × *g* for 10 min at 4°C. Crude membranes were pelleted by centrifugation at 186,007 × *g* for 45 min at 4°C using a Beckman TLA55 rotor. The resulting membrane pellets were directly processed for blue native PAGE.

### Blue native PAGE

Blue native PAGE of the crude membrane samples were performed as previously described (Zilkenat et al., 2024). Crude membrane pellets were resuspended in 1% LMNG in 1x PBS and solubilized for 1 h at 4°C with gentle shaking. The concentration of extracted membrane protein was determined using a BCA assay (Pierce™ BCA Protein Assay Kits, Thermo Fisher Scientific) following the manufacturer’s instructions. Solubilized membrane samples were mixed with the loading buffer (10×, 5% Serva Blue G in 250 mM aminocaprioic acid, 50% (w/v) glycerol) and subjected to blue native PAGE using 3-12% Bis-Tris gel (Invitrogen™ NativePAGE™). After electrophoresis, the gel was processed for immunobloting.

### Fluorescence measurements

*S.* Typhimurium strains were grown at 37°C for 5 h in LB medium with low aeration. 1 OD unit of bacterial cells was harvested by centrifugation at 5,000 × *g* for 2 min at 4°C. Pelleted cells were resuspended in 100 µl PBS and transferred to a black 96-well plate (clear bottom). Fluorescence was measured using a microplate reader (Tecan Spark). For sfGFP, excitation was set to 485 ± 10 nm and emission to 510 ± 10 nm. For mCherry, excitation was set to 587 ± 10 nm and emission to 610 ± 10 nm. Measurements were acquired in top-read mode with 100 gain, 30 flashes, and orbital shaking (2.5 mm, 216 rpm, 5 seconds) before each read. Fluorescence intensities were blank subtracted (PBS only) and the ratio between sfGFP and mCherry were obtained.

### Growth assay

*S.* Typhimurium strains were grown for 9 h at 37°C in LB medium supplemented with 0.3 M NaCl in a 96-well plate inside a humidity cassette. Bacterial growth was monitored by measuring OD_600_ every 5 min following brief shaking, using a microplate reader (Tecan Spark).

### mRNA structure predictions

All mRNA structure predictions were performed using SPOT-RNA2 with default parameters (J. Singh et al., 2021). The sequences for mRNA predictions are listed in the Supplementary Table 3, and the predicted structures were visualized using VARNA (v3-93) (Darty et al., 2009).

### AlphaFold2 predictions

Alphafold2 was used for structure predictions of SctT oligomers (Bryant et al., 2022). The list of the input sequences are listed in Supplementary Table 3.

### Statistical analysis

Statistical analysis for all graphs was performed in GraphPad Prism v10.1.1. The statistical test used in each graph is indicated in the figure legend.

## Data availability

Unmodified western blot figures and the replicates are shown in Supplementary Fig. 4.

## Acknowledgements

We thank Libera Lo Presti for critical review of the manuscript. Bioinformatic analysis was supported by the de.NBI Cloud at the University of Tübingen within the German Network for Bioinformatics Infrastructure (de.NBI).

## Author contributions

Eunjin Kim: Methodology, Investigation, Data curation, Visualization, Writing - original draft, Writing - review and editing;

Mirjam Forberger, Felix Weichel: Investigation, Funding acquisition; Claudia Paroll, Jialin Zhou, Iwan Grin: Investigation;

Samuel Wagner: Conceptualization, Methodology, Data curation, Visualization, Supervision, Writing - original draft, Writing - review and editing, Project administration, Funding acquisition.

---

**Supplementary Figure 1.**
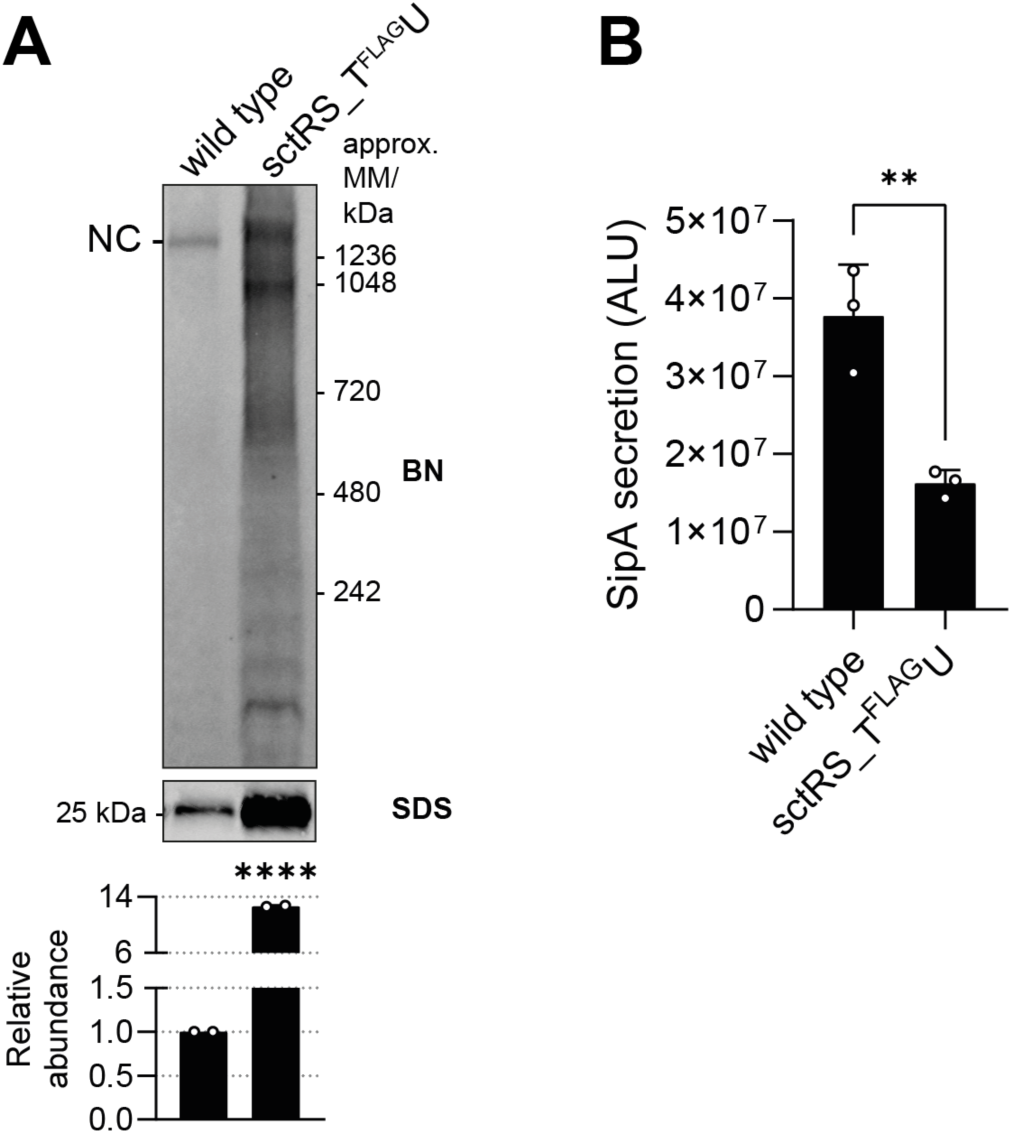
Chromosomally introduced spacer between *sctS* and *sctT* results in SctT overexpression and reduced secretion function. **A.** Equal amounts of crude membrane samples of wild-type and spacer mutant *S*. Typhimurium strains with a SipA-NL background were loaded on BN- and SDS-PAGE. The intensity of the SDS-PAGE bands was quantified using Image Studio software (Li-Cor, v5.2) and is shown as relative values to the wild-type samples, which were set to a reference value of 1. The data represent the mean of two biological replicates with the standard deviations shown as error bars. **B.** SipA-NL secretion was measured from the wild-type and spacer mutant *S*. Typhimurium strains with SipA-NL background. The P-values were calculated with an unpaired t-test, compared to the wild-type. ****, P ≤ 0.0001. ***, P ≤ 0.001, **, P ≤ 0.01.

**Supplementary Figure 2.**
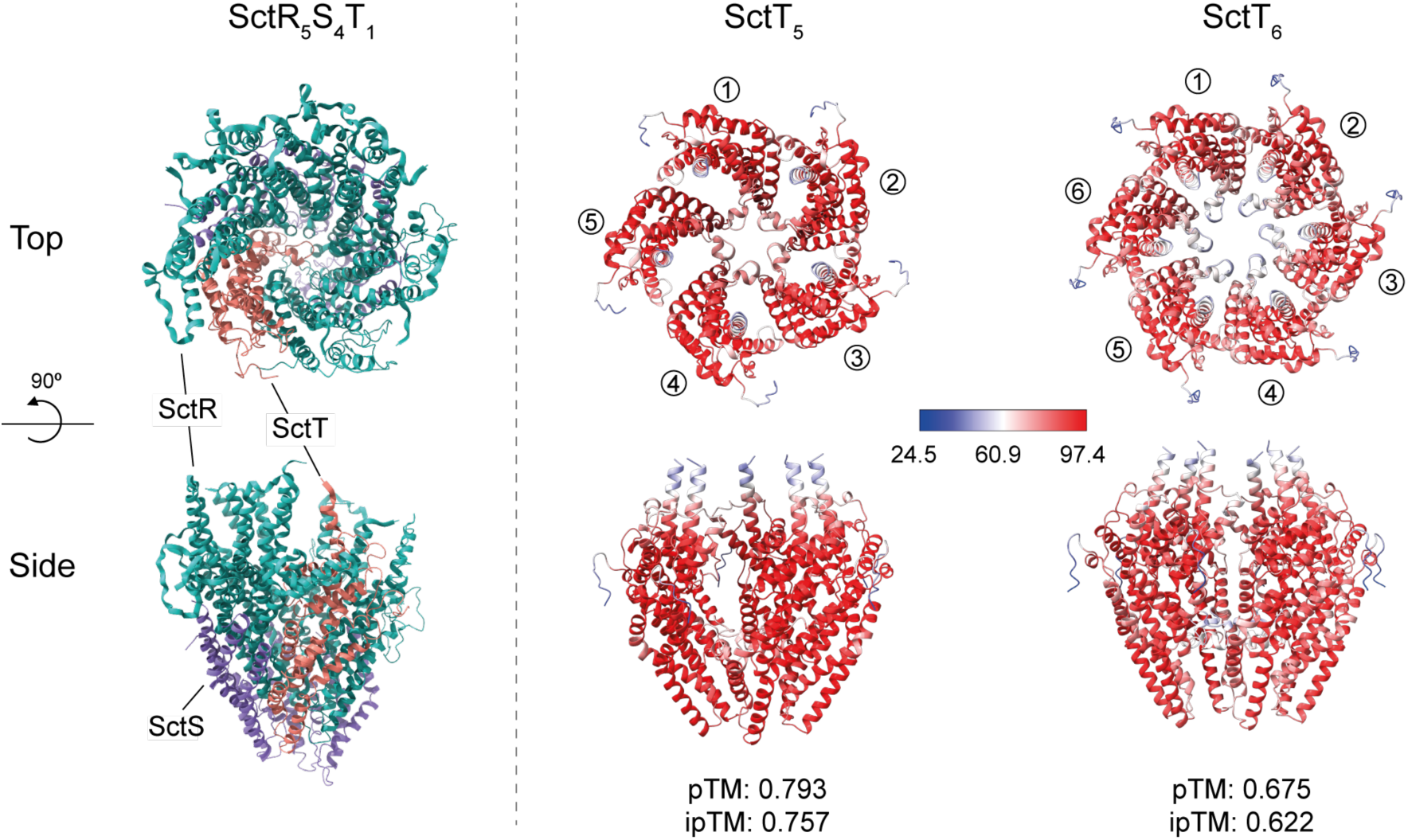
SctT multimer formation prediction by AlphaFold2. Cryo-EM structure of SctR5T1 complex (PDB 6F2D) (Kuhlen et al., 2018) and AlphaFold2 multimer predictions (Evans et al., 2021) of SctT pentamer and hexamer. In the SctR5S4T1 structure, SctR is shown in cyan, SctS in purple, and SctT in red. The confidence level of AlphaFold multimer predictions is displayed in gradients from blue (low confidence) to red (high confidence). The corresponding pTM and ipTM scores are shown below each prediction.

**Supplementary Figure 3.**
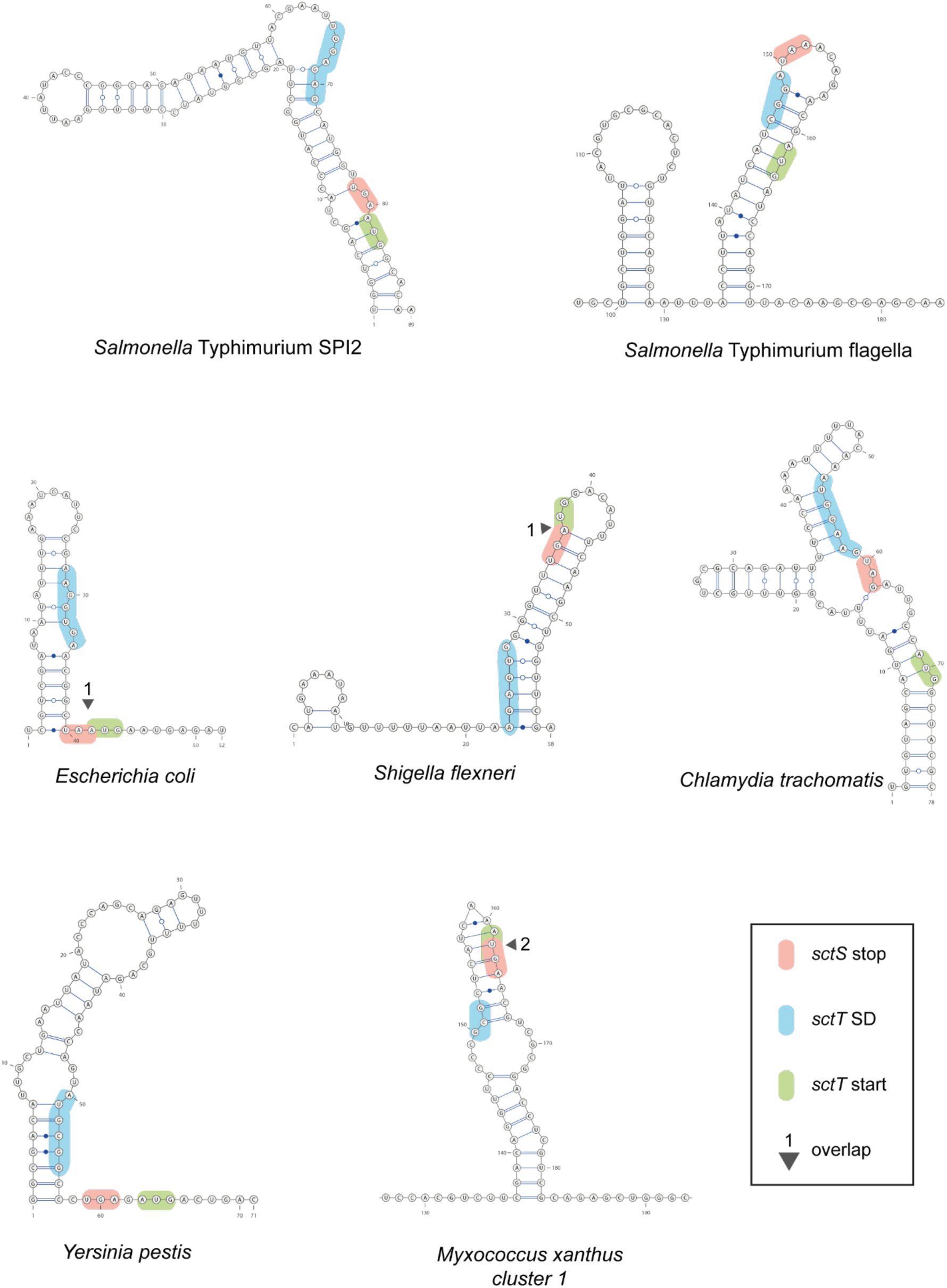
Conservation of stem-loop structure of *sctS* across many T3SS-expressing bacteria. Predicted secondary mRNA structures in the intragenic region of *sctS* and *sctT* homologs in other pathogens. The *sctS* stop codon, *sctT* start codon, and *sctT* SD sequence are shown in red, green, and blue, respectively. Overlapping *sctS* stop and *sctT* start codons are marked with a triangle and the number of overlapping bases. mRNA structure predictions were performed with SPOT-RNA2 (J. Singh et al., 2021). SPI, Salmonella pathogenicity island. SD, Shine-Dalgarno.

**Supplementary Figure 4.**
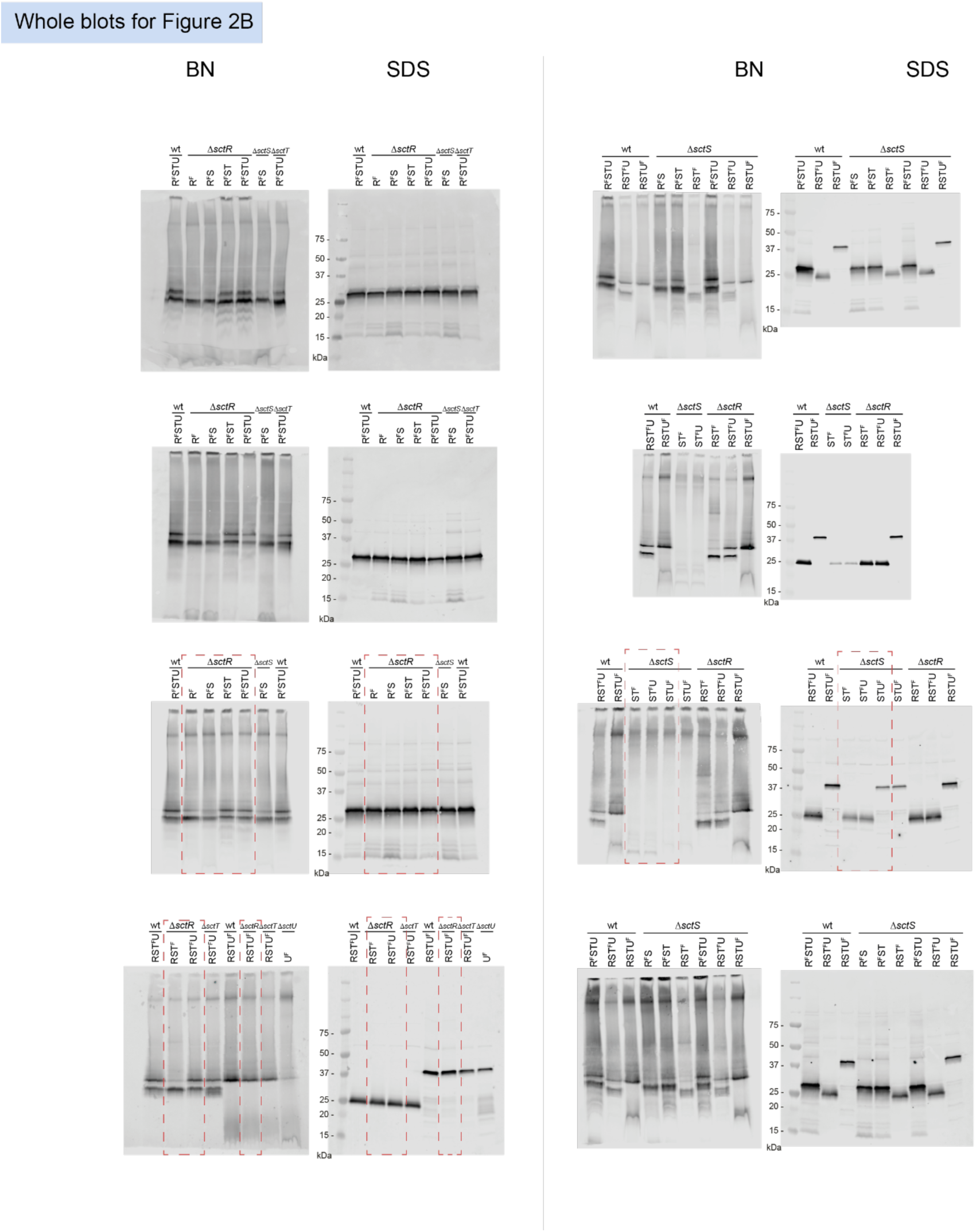
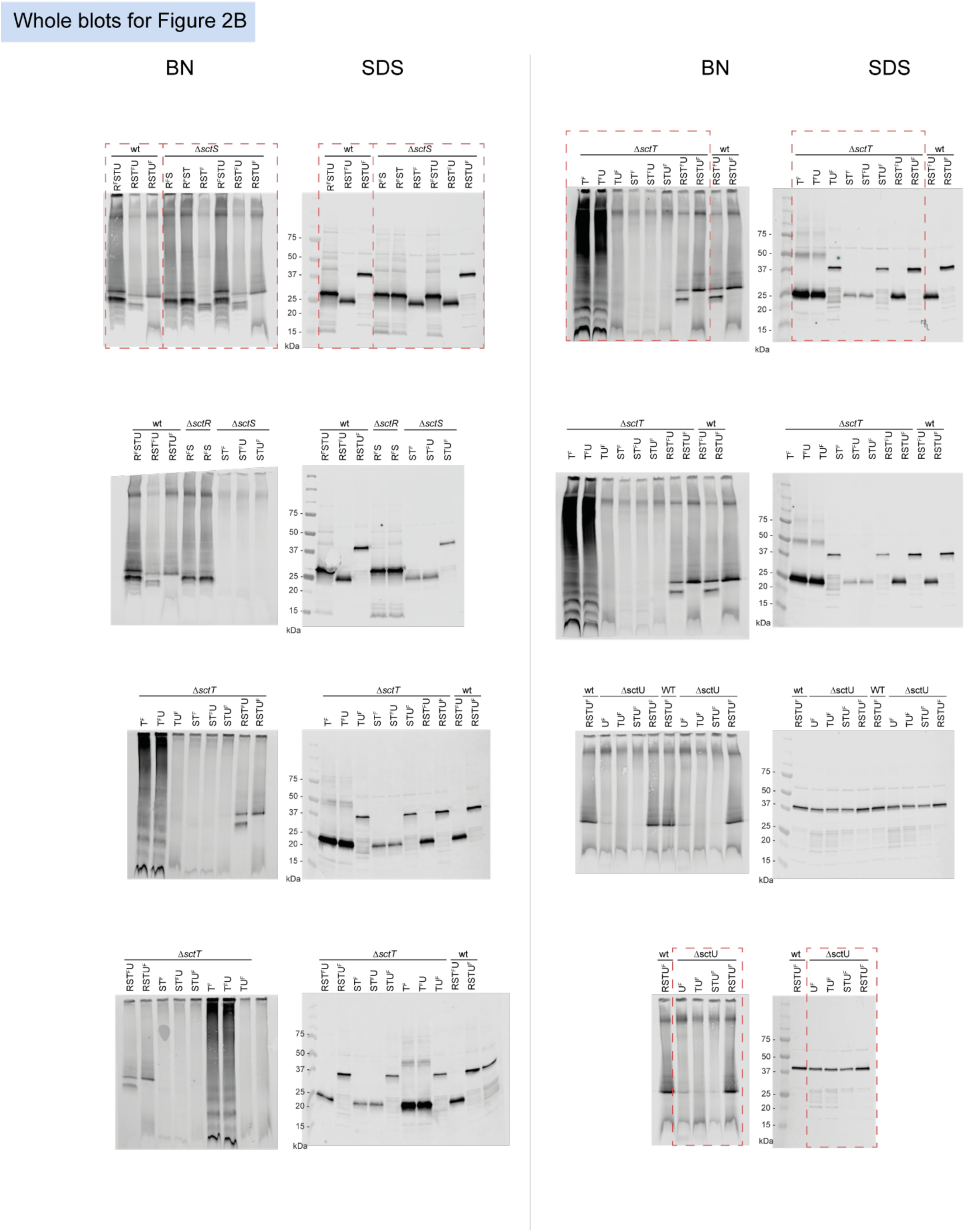
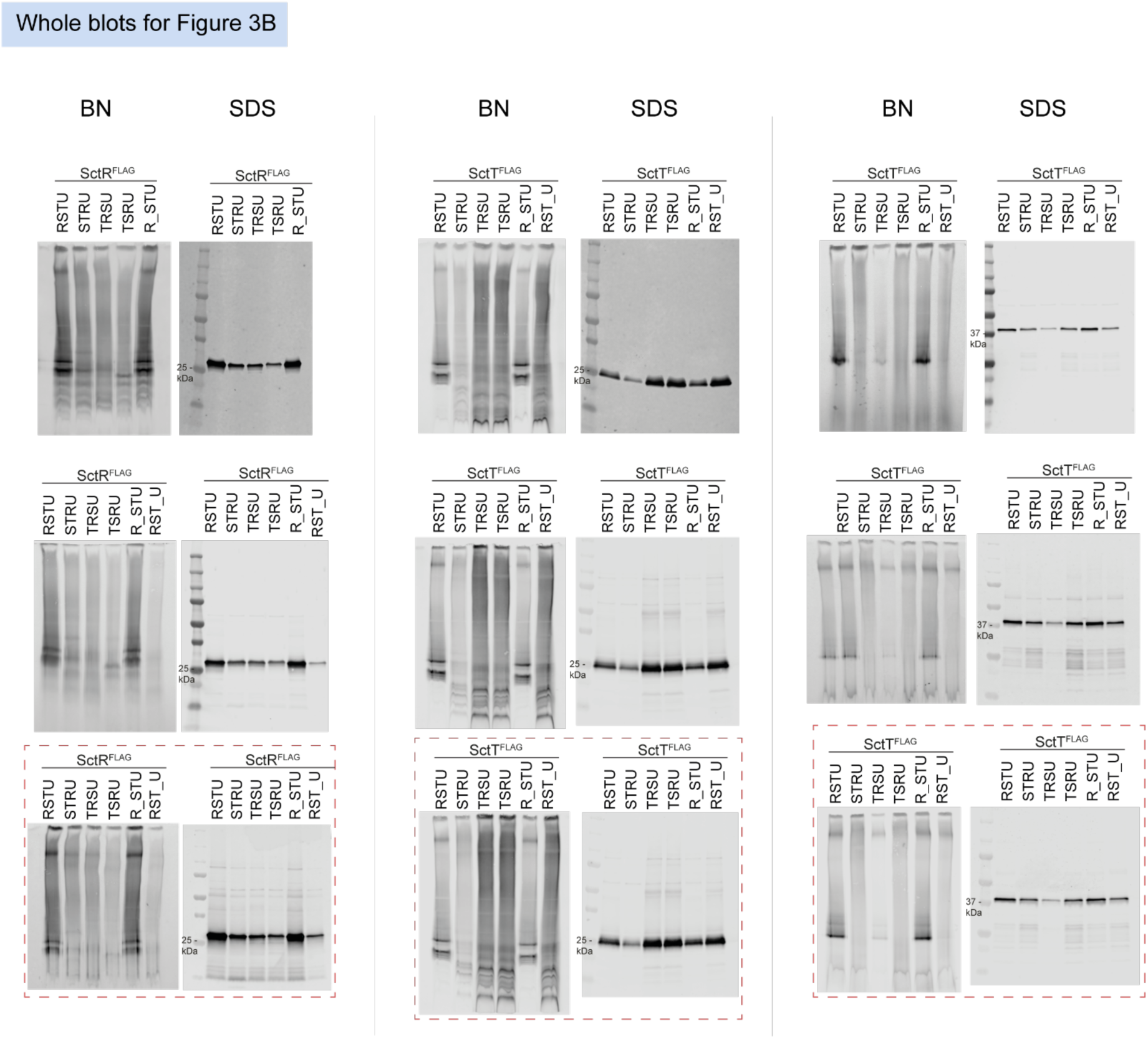
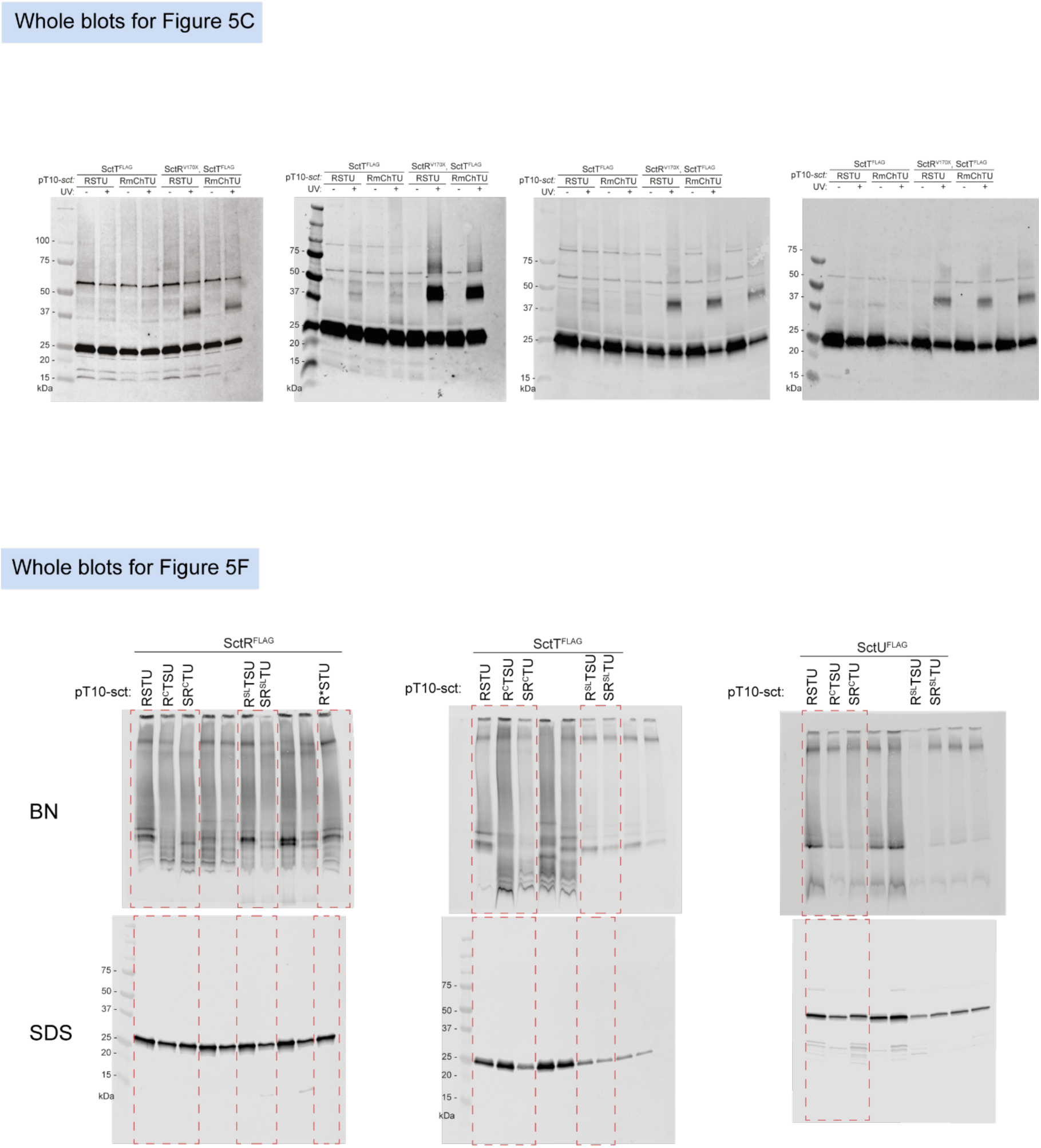
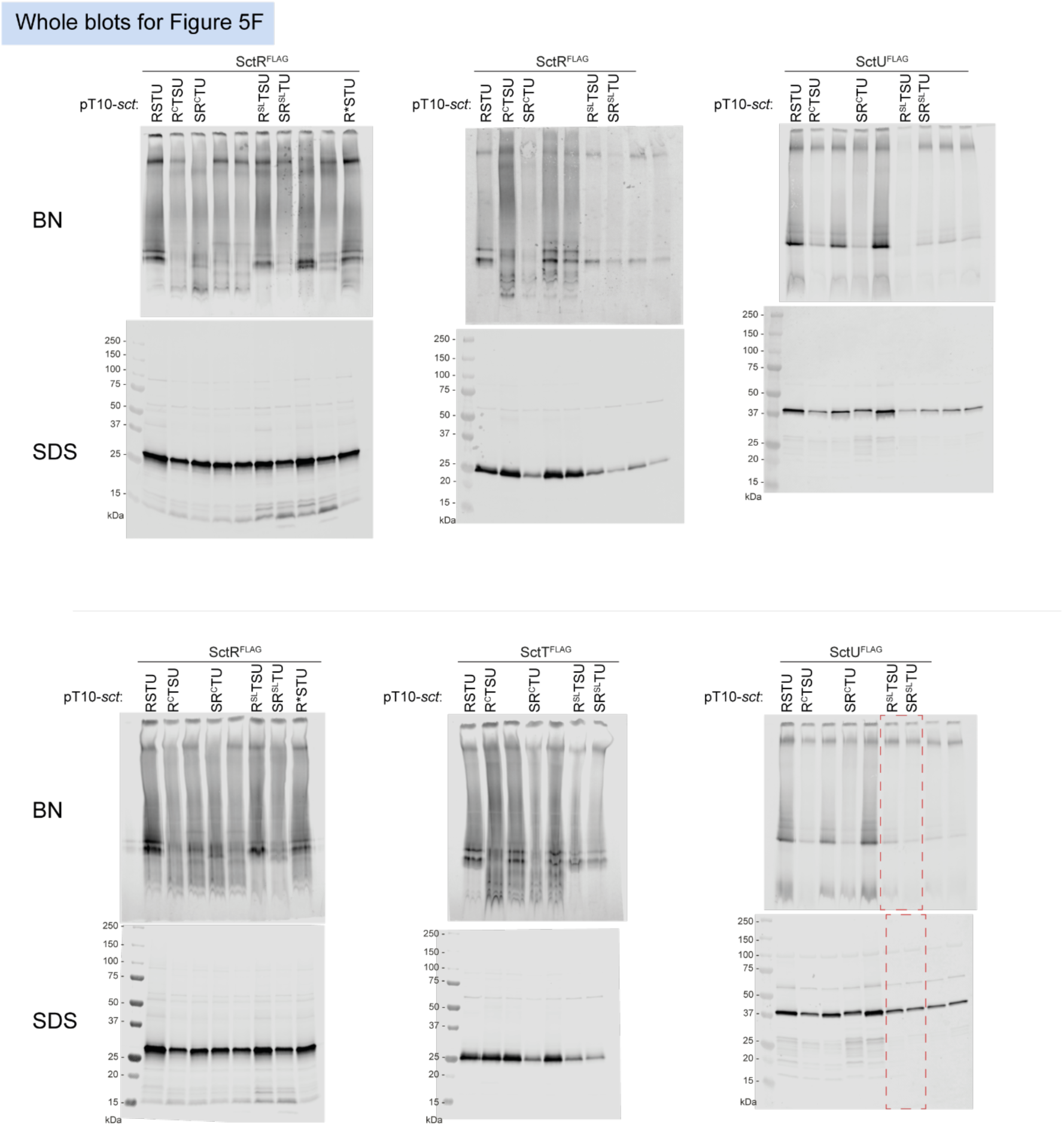
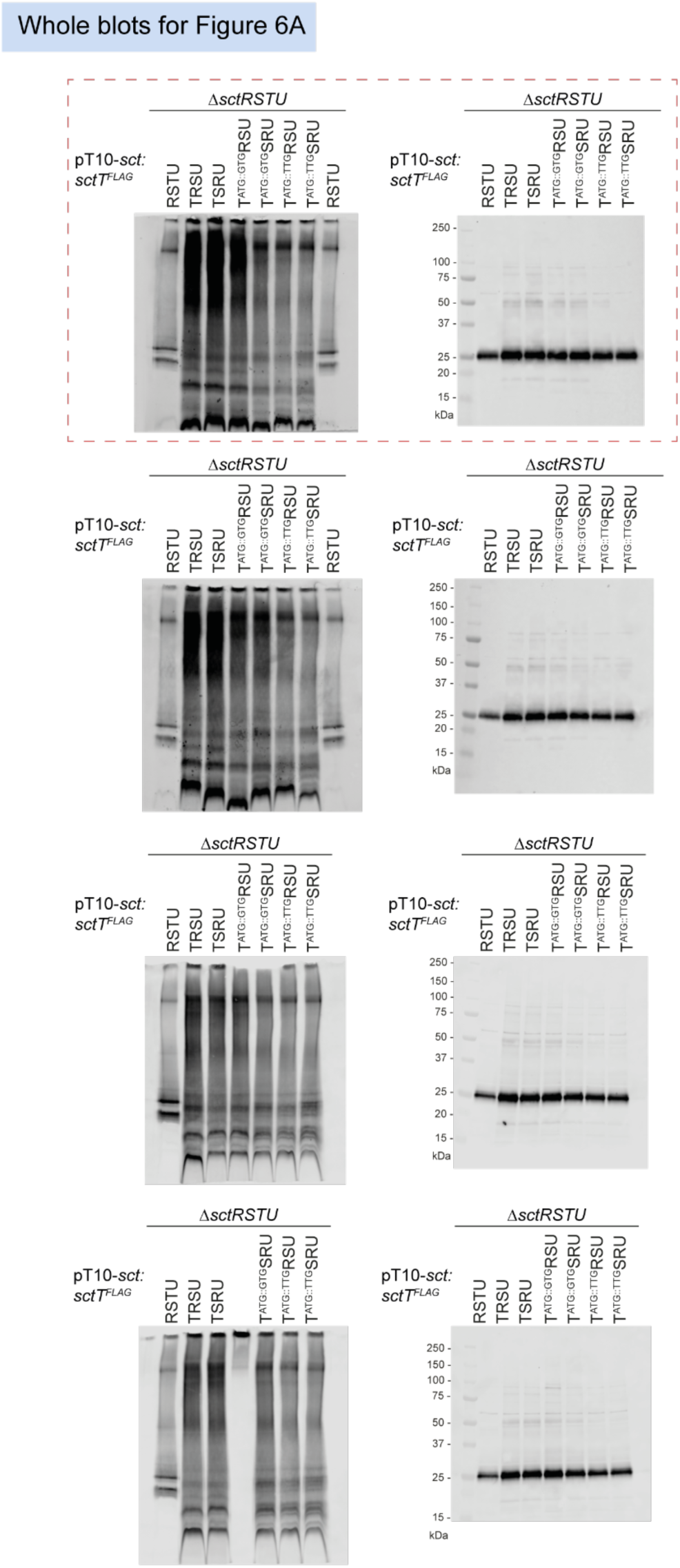
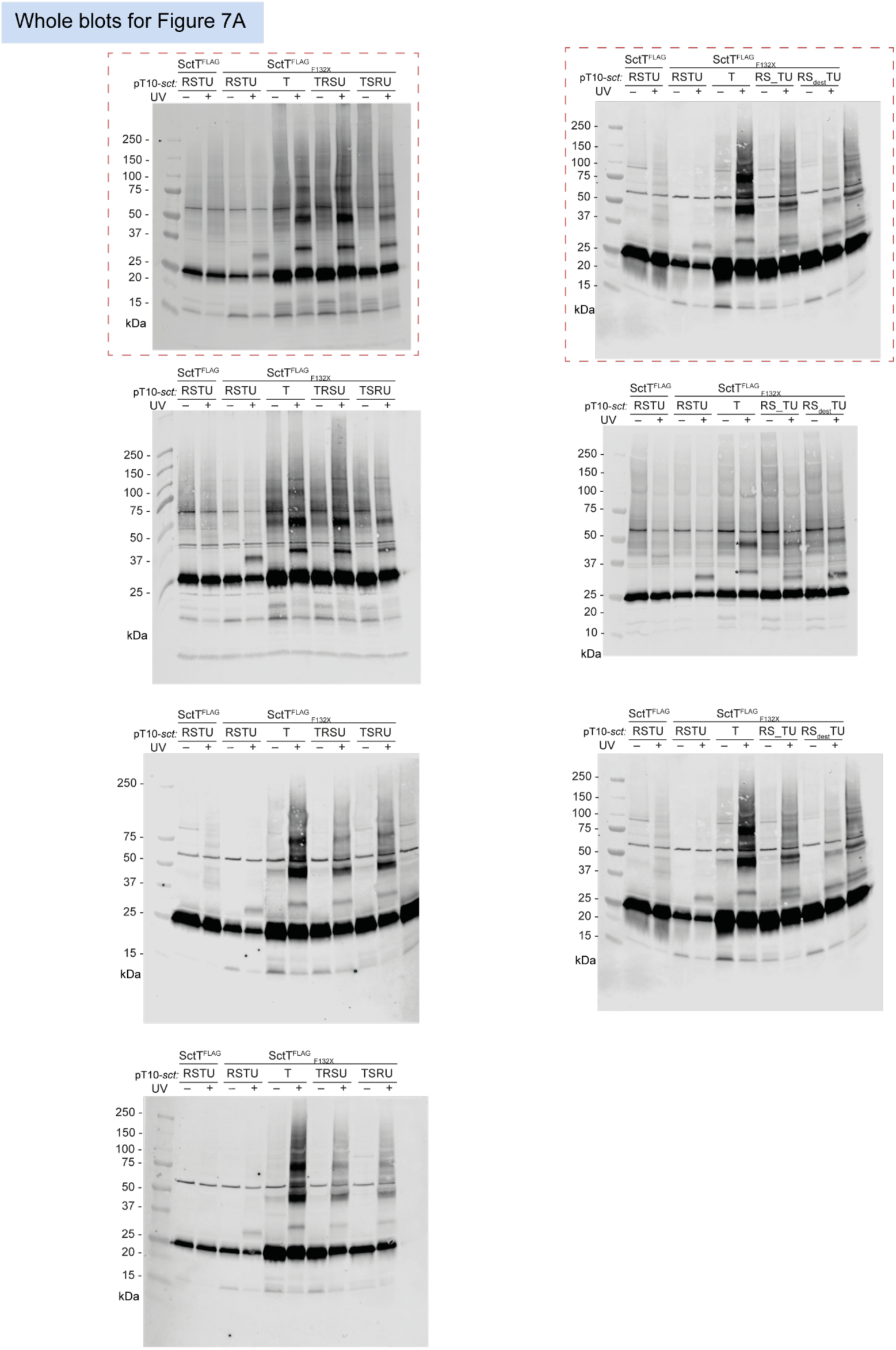
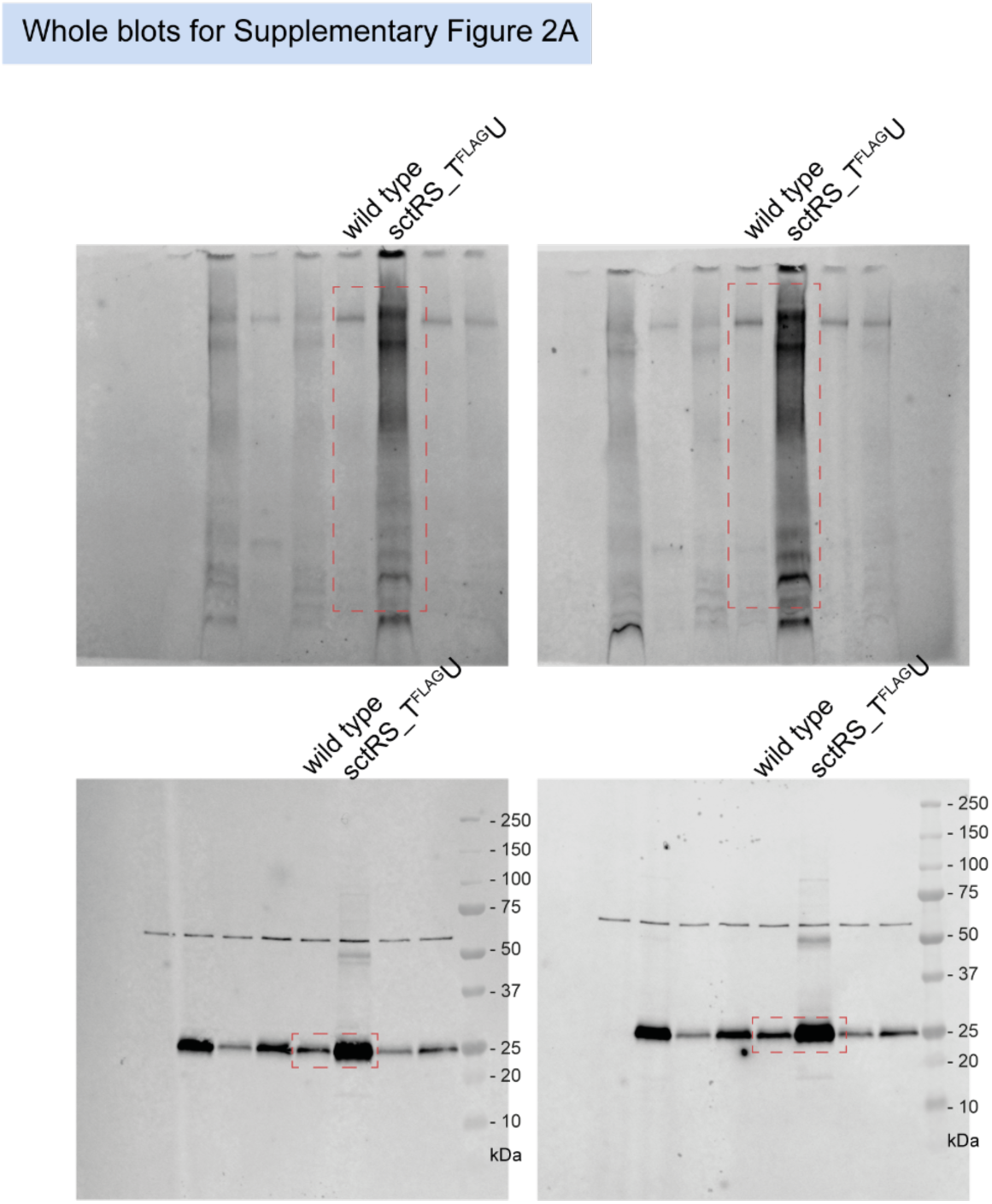
Unmodified immunoblots shown in the study. The western blots were quantified and included in the statistical analyses presented in the corresponding figures. The intensity of the SDS-PAGE bands was quantified using the Image Studio software (Li-Cor, v5.2). Blots highlighted with red dotted boxes are those shown in the main figures.

## Supplementary Information

**Supplementary Table 1. List of primers, plasmids and strains**

**Supplementary Table 2. List of hits from in silico synteny analysis**

**Supplementary Table 3. List of sequences for mRNA and protein structure predictions**

